# The biosynthesis of the odorant 2-methylisoborneol is compartmentalized inside a protein shell

**DOI:** 10.1101/2024.04.23.590730

**Authors:** Michael P. Andreas, Tobias W. Giessen

**Affiliations:** Department of Biological Chemistry, University of Michigan Medical School, Ann Arbor, MI 48109, USA

## Abstract

Terpenoids are the largest class of natural products, found across all domains of life. One of the most abundant bacterial terpenoids is the volatile odorant 2-methylisoborneol (2-MIB), partially responsible for the earthy smell of soil and musty taste of contaminated water. Many bacterial 2-MIB biosynthetic gene clusters were thought to encode a conserved transcription factor, named EshA in the model soil bacterium *Streptomyces griseus*. Here, we revise the function of EshA, now referred to as *Sg* Enc, and show that it is a Family 2B encapsulin shell protein. Using cryo-electron microscopy, we find that *Sg* Enc forms an icosahedral protein shell and encapsulates 2-methylisoborneol synthase (2-MIBS) as a cargo protein. *Sg* Enc contains a cyclic adenosine monophosphate (cAMP) binding domain (CBD)-fold insertion and a unique metal-binding domain, both displayed on the shell exterior. We show that *Sg* Enc CBDs do not bind cAMP. We find that 2-MIBS cargo loading is mediated by an N-terminal disordered cargo-loading domain and that 2-MIBS activity and *Sg* Enc shell structure are not modulated by cAMP. Our work redefines the function of EshA and establishes Family 2B encapsulins as cargo-loaded protein nanocompartments involved in secondary metabolite biosynthesis.

## Introduction

Terpenoids, also called isoprenoids, are modified terpenes and represent the largest and most structurally diverse class of natural products^1^. They play essential roles in both primary and secondary metabolism and are found across all domains of life^2,3^. They include vitamins, steroids, fragrances, plant hormones, pigments, and many compounds used as drugs^4^. While the majority of characterized terpenoids have been isolated from plants and fungi, recent genomic studies have revealed a large number of diverse terpenoid biosynthetic gene clusters encoded in bacterial genomes^2^. All terpenoids are assembled from the five-carbon (C5) prenyl diphosphate precursors dimethylallyl diphosphate (DMAPP) and isopentenyl diphosphate (IPP)^1-3,5^. Through successive additions of IPP building blocks to DMAPP, prenyl synthetases assemble linear terpenoid precursors like geranyl pyrophosphate (GPP), farnesyl pyrophosphate (FPP), and geranylgeranyl pyrophosphate (GGPP). These linear precursors can subsequently undergo a wide range of enzyme-catalyzed modifications to produce diverse and structurally complex terpenoids. Often, linear precursors are modified through carbocation-driven cyclizations catalyzed by terpene cyclases (TCs), yielding (poly)cyclic terpenoids with complex carbon scaffolds^1^. The nomenclature of terpenoids is based on the number of five-carbon units in the parent terpene with the most common bacterial terpenoids being mono- (C10), sesqui- (C15), or diterpenoids (C20)^2^.

Two of the most prevalent bacterial terpenoids are the methylated monoterpenoid 2-methylisoborneol (2-MIB) and the sesquiterpenoid geosmin, volatile odiferous compounds responsible for the earthy smell of soil and musty taste of contaminated water^2,6-8^. While the exact biological function of these volatile terpenoids is unclear, roles as insect attractants for bacterial spore dispersal^9^ and repellants against bacterial predators^10-12^ have recently been suggested. 2-MIB is widely produced by soil-dwelling Actinobacteria, Myxobacteria, and fungi, as well as by Cyanobacteria^2,6-8,13-16^. 2-MIB biosynthesis proceeds in two steps. Initially, an *S*-adenosyl methionine (SAM)- dependent methyltransferase (MT) modifies the linear C10 precursor GPP, yielding 2-methylgeranyl pyrophosphate (2-MeGPP). Next, 2-MeGPP is cyclized by the Mg^2+^- dependent class 1 terpenoid cyclase 2-methylisoborneol synthase (2-MIBS), resulting in 2-MIB formation^13,16,17^. Both proteins – MT and 2-MIBS – are generally found in co-regulated operons^13,18,19^. Commonly, a third gene encoding for a cyclic nucleotide-binding domain (CBD)-containing protein is also present in 2-MIB gene clusters^14,15,18^. This third gene was previously hypothesized to represent a transcription factor^20^ as CBDs are a common feature of bacterial transcriptional regulators and include well characterized examples like the catabolite activator protein (CAP) from *Escherichia coli*^21^. However, more recently, the CBD-containing proteins found in 2-MIB biosynthetic gene clusters have been suggested to represent Family 2B encapsulin shell proteins instead^22-24^.

Encapsulin nanocompartments (encapsulins) are the most recently discovered class of prokaryotic protein compartments able to sequester dedicated cargo enzymes^25^. Cargo encapsulation is mediated by specific targeting sequences found at the N- or C-terminus of all native cargo proteins^22,24,26,27^. Encapsulin shell proteins share the HK97 phage-like fold and likely have a viral origin^22,24,28,29^. Encapsulins self-assemble into icosahedral shells with triangulation numbers of T=1 (60 subunits), T=3 (180 subunits), or T=4 (240 subunits)^30-34^. Encapsulins have recently been categorized into four families based on sequence and genome neighborhood conservation^22^. Family 1 encapsulins are the best characterized and are involved in prokaryotic iron storage^26,31,35^, detoxification^36,37^, and stress resistance^24,38^. Family 2 encapsulins are subdivided into Family 2A and Family 2B based on the absence (2A) or presence (2B) of a CBD insertion within the encapsulin shell protein^22-24,39^. Some Family 2A encapsulins have recently been shown to encapsulate cysteine desulfurase cargo enzymes and are able to sequester large amounts of elemental sulfur^23,24,39^. Bioinformatic analyses have suggested that Family 2B encapsulins may be involved in terpenoid biosynthesis, with polyprenyl transferase- and terpene cyclase-encoding genes abundantly found in Family 2B encapsulin gene clusters^22,23^. No Family 2B encapsulin system has so far been experimentally characterized or confirmed as a protein compartmentalization system.

As mentioned above, Family 2B encapsulins have historically been annotated as CBD-containing transcription factors. This includes EshA, initially suggested to be involved in the formation of sporogenic hyphae in *Streptomyces griseus* and the regulation of antibiotic production in *Streptomyces coelicolor* A3(2)^20,40^. Purified EshA from *S. griseus* was later shown to form large multimeric complexes^41^. Surface plasmon resonance and targeted mutagenesis further suggested that the CBD of EshA interacts with cyclic adenosine monophosphate (cAMP)^42^. When these early studies were being carried out, encapsulins had not yet been discovered and no further characterization of EshA has been reported since.

Here, we revise the function of EshA from *S. griseus* and establish it as a Family 2B encapsulin shell protein that sequesters 2-MIBS as its cargo enzyme. Using cryogenic electron microscopy (cryo-EM), we report the structure of a Family 2B encapsulin shell and highlight the presence of a CBD insertion domain and a previously not recognized metal-binding domain within the Family 2B shell protein. We show that 2-MIBS cargo-loading is mediated by a flexible proline-rich N-terminal cargo-loading domain (CLD). We find that 2-MIBS encapsulation does not significantly affect enzyme activity or product distribution and that the presence of cAMP does similarly not modulate 2-MIBS activity. Detailed binding studies further suggest that CBDs in Family 2B encapsulin shell proteins do not bind cAMP. Our study establishes Family 2B encapsulins as protein nanocompartments with a unique shell architecture and highlights that the biosynthesis of 2-MIB – one of the most prevalent terpenoid natural products – takes place inside an encapsulin shell in the majority of bacteria. Our work lays the groundwork for further elucidating the structure and function of Family 2B encapsulin systems.

## Results

### The distribution and diversity of 2-MIB biosynthetic gene clusters in Actinobacteria

In this study, we focus on the 2-MIB biosynthetic gene cluster found in *S. griseus*, encoding *Sg* Enc (formerly EshA), 2-MIBS, and a methyltransferase (MT) (Fig. 1a). 2-MIB biosynthetic gene clusters are ubiquitous in Actinobacteria, most notably in soil-dwelling *Streptomyces,* which are renowned for their ability to produce diverse secondary metabolites, including volatile terpenoids^2,6,8^. 2-MIBS catalyzes the cyclization of methylated 2-methylgeranyl pyrophosphate (2-MeGPP) and was the first terpene cyclase observed to utilize a non-canonical isoprenoid substrate (Fig. 1b)^13,16,17^. Further, a SAM-dependent MT, responsible for converting GPP to 2-MeGPP, is also widely conserved across 2-MIBS-encoding gene clusters^13,18,19^. Genes encoding for Family 2B encapsulin shell proteins are present in many 2-MIBS-encoding operons, including in *S. griseus*^22^. Most 2-MIBS enzymes contain two main domains, a disordered ca. 120 residue N-terminal domain rich in proline, glycine, and alanine and a C-terminal terpene cyclase domain^17^. The disordered N-terminal domains have recently been proposed to function as cargo loading domains (CLD) in Family 2B encapsulin systems^22^. Similar N-terminal CLDs are found in Family 2A cysteine desulfurase cargo proteins and have already been confirmed as essential for cargo encapsulation^22,23,39,43^.

**Fig. 1.**
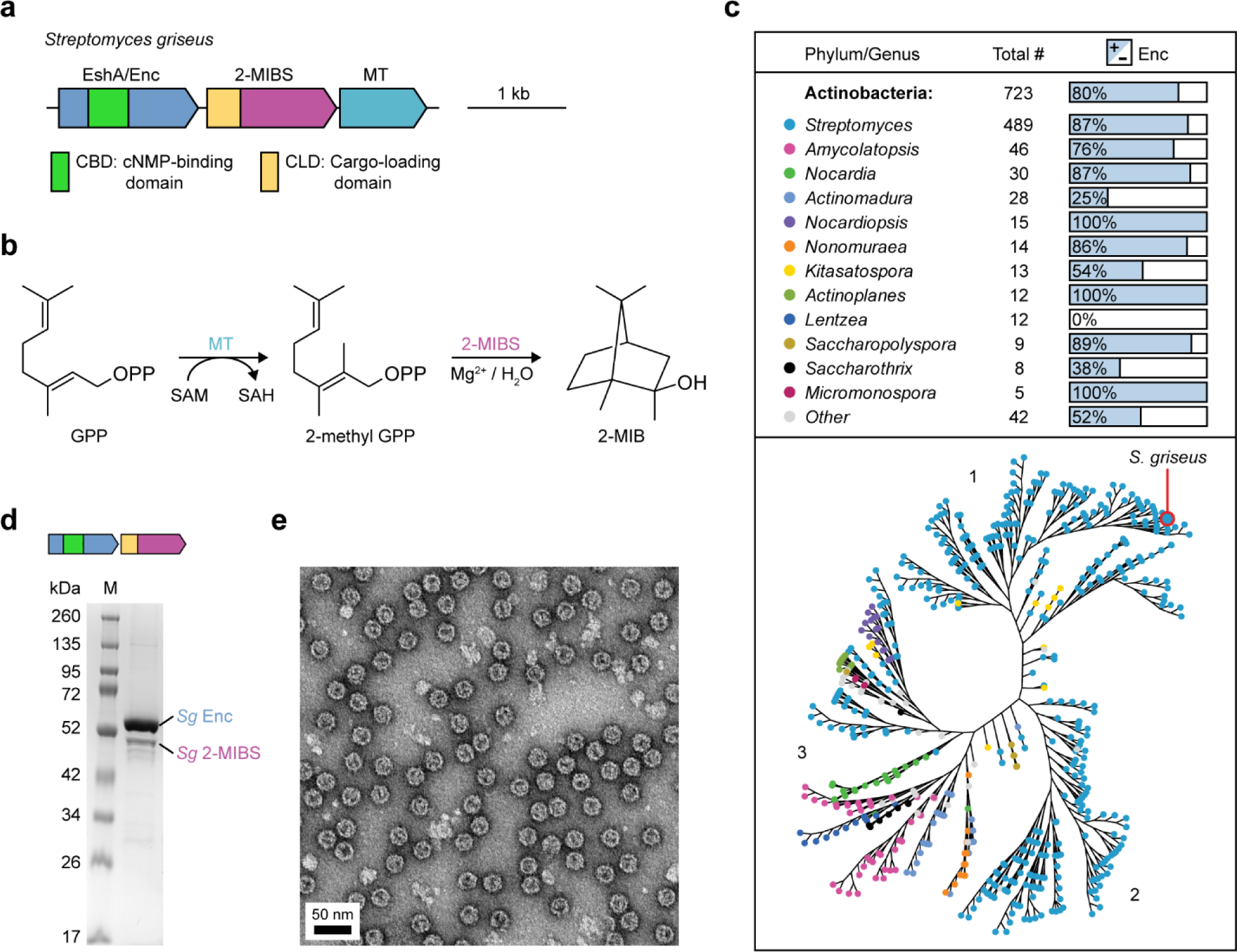
2-methylisoborneol biosynthetic gene clusters encoding a Family 2B encapsulin shell protein and its 2-MIBS enzyme cargo are widespread in Actinobacteria. **a,** The 2-methylisoborneol (2-MIB) biosynthetic gene cluster from *Streptomyces griseus* (*Sg*). The operon encodes a Family 2B encapsulin shell protein (EshA/Enc), 2-methylisoborneol synthase (2-MIBS), and a methyltransferase (MT). kb: kilobases. **b,** Reaction scheme for the biosynthesis of 2-MIB. Geranyl pyrophosphate (GPP) is methylated by the MT, yielding 2-methyl GPP, followed by 2-methyl GPP cyclization catalyzed by 2-MIBS resulting in 2-MIB formation. SAM: *S*-adenosyl methionine. SAH: *S*-adenosyl homocysteine. **c,** Phylogenetic distribution and encapsulin prevalence in actinobacterial 2-MIB biosynthetic gene clusters (top). Phylogenetic analysis of 723 2-MIBSs found in actinobacterial 2-MIB biosynthetic operons (bottom). Branch numbers (1 to 3) are shown. Nodes are colored by bacterial genus. **d,** SDS-PAGE analysis of purified cargo-loaded *Sg* Enc shells resulting from heterologous expression in *Streptomyces coelicolor* A3(2). **e,** Negative stain transmission electron microscopy (TEM) images of purified 2-MIBS-loaded *Sg* Enc.

Computational searches and genome neighborhood analysis yielded 723 2-MIB biosynthetic gene clusters in Actinobacteria, with the majority (67.6%) found in the genus *Streptomyces* (Fig. 1c). Family 2B encapsulin-encoding genes were identified in 80% of the operons. The remaining 20% of organisms did not encode any Family 2B encapsulins, even outside the 2-MIB biosynthetic locus, in their genomes. These findings suggest that Family 2B encapsulin association is a common but not essential feature of bacterial 2-MIB biosynthesis.

Phylogenetic analysis of actinobacterial 2-MIBSs yielded three main branches reflecting sequence similarity (Fig. 1c). Branches one and two contained primarily *Streptomyces*-encoded 2-MIBSs. Sequence alignments of these 2-MIBSs showed poor sequence conservation within CLDs (branch 1 CLDs: 22.5% average pairwise identity; branch 2 CLDs: 41.3% average pairwise identity), but high conservation of the terpen cyclase (TC) domains (branch 1 TCs: 79.9% average pairwise identity; branch 2 TCs: 80.8% average pairwise identity). Branch three contained sequences from 36 genera including *Streptomyces, Actinomadura, Nocardia,* and *Amycolatopsis.* Consequently, both the CLD and TC domains within branch three were significantly more divergent than sequences within branches one and two (branch 3 CLDs: 14.9% average pairwise identity; branch 3 TCs: 48.7% average pairwise identity). Sequence similarity network analysis yielded similar results, with two large clusters corresponding to branches one and two, and branch three broken into several smaller clusters (Supplementary Fig. 1, a and b). Sequence alignments of the 2-MIBSs within each cluster revealed short, conserved motifs within CLDs, with most clusters containing a GPxGLGT consensus motif (Supplementary Fig. 1c), which was also highlighted in previous bioinformatic analyses of putative Family 2 encapsulin cargos^22,23^. Only cluster 2 exhibited a divergent CLD consensus motif (LPGPP) instead (Supplementary Fig. 1c). Further analysis of the CLDs within each cluster showed significant variability in length, with cluster 2 containing the shortest CLDs (31 ± 12 residues), and many CLDs exceeding 200 residues in length and containing multiple repeats of the abovementioned consensus motifs (Supplementary Fig. 1c).

### EshA is a Family 2B encapsulin with 2-MIBS as its cargo protein

*Sg* Enc (formerly EshA) has previously been annotated as a CBD-containing transcription factor^41^ and has more recently been suggested to represent a Family 2B encapsulin shell protein^22^. To investigate if *Sg* Enc is indeed able to form cargo-loaded protein nanocompartments, we heterologously expressed a two-gene operon encoding *Sg* Enc and *Sg* 2-MIBS (*Sg* Enc_2-MIBS) in *Streptomyces coelicolor* A3(2). After ammonium sulfate precipitation and size-exclusion chromatography, clear co-purification of both proteins could be observed (Fig. 1d). Negative-stain transmission electron microscopy (TEM) confirmed the formation of protein shells with a diameter of ca. 28 nm, consistent with a T=1 icosahedral assembly as seen in other T=1 encapsulin shells (Fig. 1e)^23,24,30,39^. Additional densities in the interior of many shells could be observed, likely representing encapsulated 2-MIBS. Regularly spaced protrusions localized on the surface of shells could also be visualized via TEM, which may represent externally displayed CBDs.

### Single particle cryo-EM analysis of cargo-loaded *Sg* Enc reveals a unique shell architecture

To investigate the molecular structure and organization of this Family 2B encapsulin system, we carried out single-particle cryo-EM analysis of both the cargo-loaded shell (*Sg* Enc_2-MIBS) and the empty shell without cargo (*Sg* Enc). Icosahedral (I) refinement of *Sg* Enc_2-MIBS and *Sg* Enc yielded cryo-EM maps with global resolutions of 3.02 Å and 2.71 Å (Supplementary Fig. 2 and 3). Consistent with our negative stain TEM analysis, the encapsulin shell was found to possess a diameter of 29.4 nm and consist of 60 protomers of the *Sg* Enc shell protein arranged with T=1 icosahedral symmetry (Fig. 2a). CBDs were clearly visible on the exterior of the shell, albeit at significantly lower local resolution than the HK97-like shell-forming domain of *Sg* Enc (Supplementary Fig. 2d and 3d), arranged around all two-fold axes of symmetry (Fig. 2a). Additional non-*Sg* Enc density was observed lining the shell interior at the three-fold axes of symmetry for the cargo-loaded system (*Sg* Enc_2-MIBS), reminiscent of CLD density previously observed in cargo-loaded *Synechococcus elongatus* and *Acinetobacter baumannii* Family 2A encapsulin shells (Fig 2a)^23,39^. This extra density was not observed in the empty *Sg* Enc shell, further suggesting that it represents part of the 2-MIBS CLD. Apart from the surface-exposed CBDs, the overall architecture of the HK97-like shell is similar to characterized Family 2A shells, including the characteristic turret-like vertices formed at the five-fold symmetry axes (Fig. 2b and Supplementary Fig. 4a,b). In the *Sg* Enc shell, the pentameric vertices are noticeably more flattened compared to Family 2A shells due to a shorter C-terminal extension in the *Sg* Enc protomer. Small pores, ca. 4 Å and 2 Å, can be found at the five- and three-fold axes of symmetry, respectively, comparable in size to pores found in Family 2A encapsulin shells (Fig. 2b,c and Supplementary Fig. 4c). Along all two-fold symmetry axes, the *Sg* Enc shell exhibits large, elongated pores, approximately 60 x 25 Å in dimension (Fig. 2d), located beneath the externally displayed CBDs. No encapsulin shell has previously been observed to possess such large pores. In particular, pores at the two-fold axes of symmetry are usually small or non-existent in Family 1 or Family 2A encapsulin shells. Specifically, in the *S. elongatus* and *A. baumannii* Family 2A shells, the N-arm of the HK97-like domain is threaded along the two-fold axes, resulting in essentially closed two-fold pores (Supplementary Fig. 4d)^23,39^. We were able to confidently model most of the *Sg* Enc HK97-like domain, which maintained a nearly identical fold compared to Family 2A encapsulin protomers (Fig. 2e and Supplementary Fig. 4e). However, the first 39 residues of the *Sg* Enc N-arm are unresolved, resulting in the observed elongated open two-fold pores (Supplementary Fig. 5a). Closer analysis of our cryo-EM maps revealed a segment of poorly resolved density positioned adjacent to the CBDs near the N-termini (Supplementary Fig. 5b). This density may correspond to parts of the unmodelled N-arm segment of *Sg* Enc or to an unresolved loop connecting the CBDs to the E-loops of the HK97-like domains (Supplementary Fig. 5a).

**Fig. 2.**
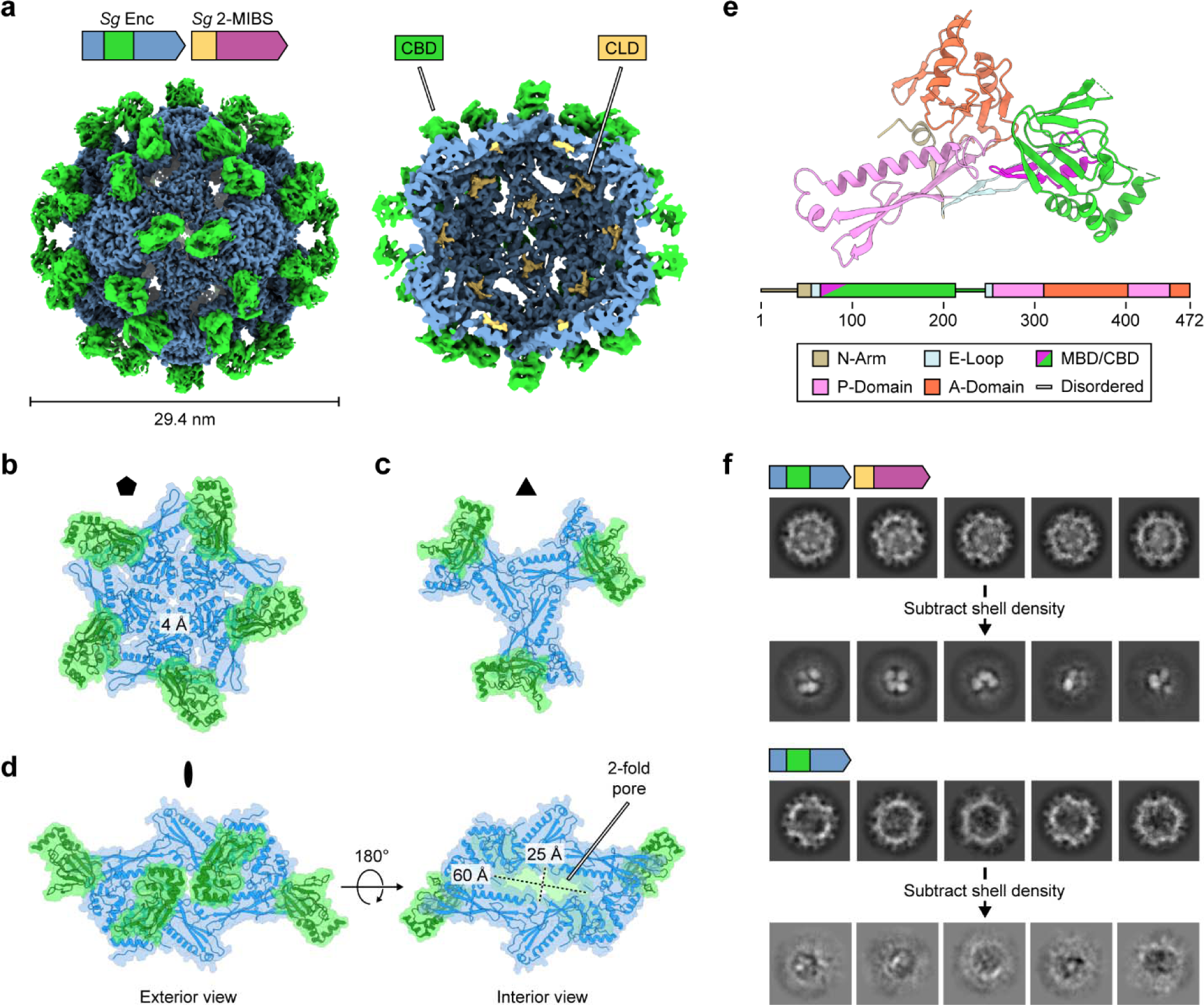
Cryo-EM analysis of 2-MIBS-loaded *Sg* Enc. **a**, Cryo-EM map of the cargo-loaded *Sg* Enc shell viewed from the exterior down the two-fold axis of symmetry (left). Cut-away view shown on the right. Density corresponding to the HK97-like shell-forming domain is shown in blue. Externally displayed CBDs are shown in green. Internalized 2-MIBS CLD densities shown in yellow. **b**, Exterior view of a *Sg* Enc pentamer down the five-fold axis of symmetry highlighting the five-fold pore. **c**, Exterior view of a *Sg* Enc trimer down the three-fold axis of symmetry. **d**, Exterior (left) and interior (right) view down the two-fold axis of symmetry highlighting the large, elongated two-fold pore. When viewed from the exterior, the CBDs partially occlude the two-fold pore. **e**, Structural model of the *Sg* Enc shell protomer with characteristic HK97-like domain elements and the CBD highlighted with different colors. MBD: metal-binding domain. **f**, 2D class averages of shell-subtracted 2-MIBS-loaded *Sg* Enc particles (top). 2D class averages of shell-subtracted empty *Sg* Enc particles (bottom). While discrete internalized cargo puncta are visible for the cargo-loaded sample, no puncta can be seen in empty shells.

The shells resulting from both I and C1 refinements of cargo-loaded *Sg* Enc_2-MIBS exhibited a significant amount of internal noise which we attribute to encapsulated but fairly mobile 2-MIBS cargo. Shell density subtraction followed by 2D classification of shell-subtracted particles produced classes with three to four distinct puncta in 99% of particles. This observation suggests that approximately three to four dimers of 2-MIBS, previously shown to form dimers^17^, are contained within each shell of *Sg* Enc_2-MIBS. In contrast, no internal densities were observed in shell-subtracted particles of the *Sg* Enc shell without cargo (Supplementary Fig. 2 and Supplementary Fig. 6). Contrary to the *A. baumannii* Family 2A cargo-loaded encapsulin, where low resolution lobes of density corresponding to cysteine desulfurase cargo could be clearly observed in the interior of the shell^39^, none of our reconstructions were able to resolve the interior orientations of the 2-MIBS cargo. This suggests that 2-MIBS may not be as spatially confined as the cysteine desulfurase cargo in Family 2A encapsulins, exhibiting substantial flexibility instead.

### The *Sg* Enc shell protein contains unique CBD and metal-binding insertion domains

While we were able to confidently dock an AlphaFold model of the CBD into the cryo-EM density resulting from I refinement of the *Sg* Enc shell, the CBD densities exhibited noticeably worse resolution than the HK97-like domain of the shell. Masked 3D classification of the CBD densities surrounding the two-fold axis of symmetry, followed by homogeneous C1 reconstruction of the best class, significantly improved the quality of the CBD density (Supplementary Fig. 7 and 8), now enabling confident CBD model building. Similar to other CBDs, the *Sg* Enc CBD is composed of an eight stranded β-barrel domain and an α-helical subdomain composed of three short α-helices (Fig. 3a). This overall fold, and the β-barrel in particular, is strongly conserved as the primary cAMP-binding site in characterized cAMP-binding proteins such as cAMP-dependent protein kinase A, ion-gated channels, and cAMP-responsive transcription factors, most notably the *E. coli* catabolite activator protein (*Ec* CAP) (Fig. 3b)^44,45^. In many cAMP-regulated proteins, binding of cAMP to the β-barrel domain results in a conformational change that is propagated throughout the protein or complex to regulate the activity of a distal functional domain. In the dimeric *Ec* CAP for example, cAMP binding induces a conformational change that stabilizes a central coiled-coil helical domain and re-orients the DNA-binding domains to enable stable interaction with double-stranded DNA^46,47^. Instead of an extended α-helix able to form a coiled-coil, as found in *Ec* CAP, the C-terminus of monomeric *Sg* Enc CBD contains a long disordered loop that is unresolved in our cryo-EM density (Supplementary Fig. 5a). This loop acts as a linker between the CBD and the HK97-like domain of *Sg* Enc and may also play a role in ligand binding or modulating the conformation of the CBD in analogy to the extended α-helices observed in other CBD containing proteins.

**Fig. 3.**
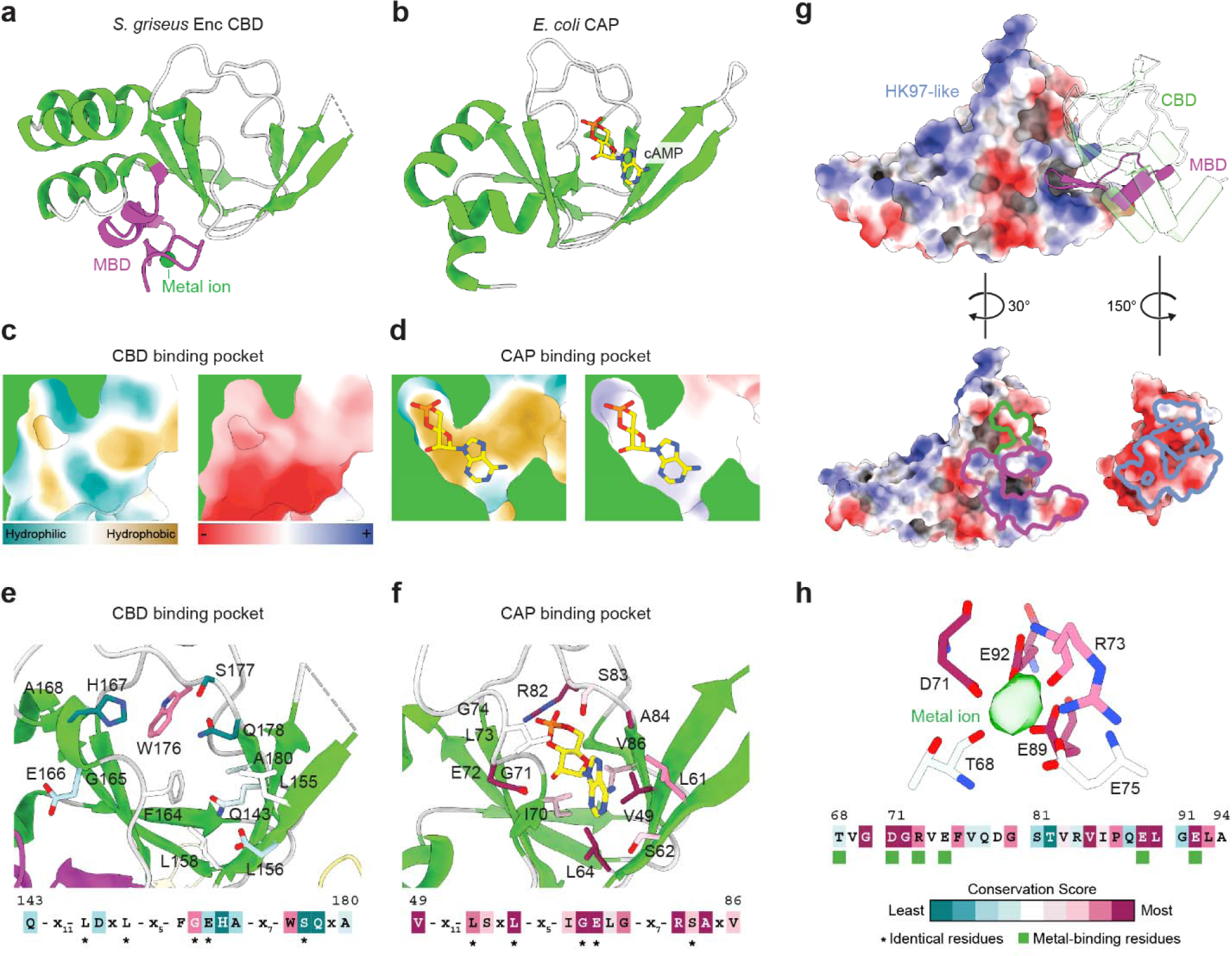
Structural analysis of the *Sg* Enc CBD. **a**, An overview of the structure of the *Sg* Enc CBD. The metal-binding subdomain (MDB) is shown in purple and the remainder of the CBD is shown in green and light grey. **b**, The *E. coli* (*Ec*) CAP CBD (residues 1 to 116) bound to cAMP (PDB ID: 1G6N)^21^ colored analogously to panel **a**. cAMP is shown as yellow sticks. **c**, Surface representations of the *Sg* Enc CBD binding pocket colored according to hydrophobicity (left) and surface charge (right). **d**, Surface representations of the *Ec* CAP CBD binding pocket colored according to hydrophobicity (left) and surface charge (right). Bound cAMP is shown as yellow sticks. **e**, Key *Sg* Enc CBD residues forming the CBD binding pocket. Individual residues are colored by conservation as predicted by ConSurf analysis^48-50^. ConSurf conservation score key shown in panel **h**. **f**, Key *Ec* CAP CBD residues in the CBD binding pocket involved in cAMP binding. Individual residues are colored by conservation as predicted by ConSurf analysis. **g**, The CBD interacts with the HK97-domain of the *Sg* Enc protomer predominantly via electrostatic interactions mediated by the MBD. The interaction interfaces are outlined on the HK97-domain surface (left) for the CBD (green) and the MBD (purple). Conversely, interaction interfaces of the HK97-domain with the CBD and MBD are outlined in blue (right). **h**, The MBD is a highly conserved feature of Family 2B encapsulin shell proteins. Side chains and backbone atoms coordinating the metal ion are colored according to sequence conservation as predicted by ConSurf analysis. Extracted cryo-EM metal ion density is shown in green.

No ligands were observed bound to the CBDs for either the cargo-loaded *Sg* Enc_2-MIBS or shell-only *Sg* Enc CBD samples, despite the samples not having been subjected to dialysis or ion exchange chromatography. Closer analysis of the β-barrel binding pocket shows that in *Sg* Enc CBD, a dramatically different hydrophobicity and surface charge pattern can be observed than in *Ec* CAP (Fig. 3, c and d). Furthermore, the usually highly conserved cAMP-interacting residues found in CBDs, including *Ec* CAP, are poorly conserved in *Sg* Enc CBD (Fig. 3, d and e). In particular, a highly conserved arginine residue located at the back of the binding pocket – R82 in *Ec* CAP – is replaced with a tryptophan residue – W176 in *Sg* Enc – in essentially all Family 2B encapsulin CBDs. In canonical cAMP-binding CBDs, the conserved arginine forms part of the phosphate binding cassette (PBC), essential for cAMP binding, and directly coordinates the cyclic phosphate of cAMP^44,45^. Mutations to R82 of *Ec* CAP generally result in a dramatic loss of cAMP binding^51-53^. CBD-like domains lacking this arginine are not responsive to cAMP^45^, and have been shown to instead bind a diverse array of other ligands such as heme in the CO-responsive transcription factor CooA^54^, chlorophenolacetic acid in the transcriptional regulator CprK^55^, 2-oxoglutarate in the transcription factor NtcA^56^, and even peptides in the case of KCNH voltage-gated potassium ion channels^57-59^. Because *Sg* Enc, and nearly all identified Family 2B encapsulins, lack this conserved arginine, it is possible that Family 2B encapsulin-associated CBDs may not bind cAMP but alternative ligands instead.

Closer analysis of the N-terminal section of the *Sg* Enc CBD revealed a 30 residue motif, which we will refer to as the metal-binding domain (MBD), composed of two anti-parallel β-strands and a short α-helix that form a metal binding site, not found in other characterized CBDs (Fig. 3a). This motif appears to be unique to Family 2B encapsulins and similar motifs could not be readily identified in known protein structures via structural homology searches using DALI, HHPRED, or FoldSeek^60-63^. The negatively charged MBD forms an interface with a positively charged surface patch on the exterior of the HK97-like *Sg* Enc domain, which may function to stabilize the orientation of the CBD (Fig. 3g). The CBD, in comparison, maintains a much smaller interface with the HK97-like domain, primarily mediated by hydrophobic interactions. An unknown metal ion, likely Mg^2+^ or Ca^2+^ based on the observed coordination pattern, is bound by conserved residues, primarily glutamates and aspartates, within the metal binding site of the MBD (Fig. 3, a and h), suggesting that metal ion binding is a widespread feature of the MBD throughout Family 2B encapsulins.

### 2-MIBS encapsulation is mediated by a disordered cargo-loading domain

As mentioned above, additional cryo-EM density, likely belonging to a fragment of the 2-MIBS CLD, was observed in the cargo-loaded *Sg* Enc shell interior located at the three-fold axes of symmetry. In I refinements, these additional densities appeared symmetry-averaged and could not be used for model building due to their limited resolution and quality. Asymmetric refinements yielded CLD densities which appeared to partially break the three-fold symmetry (Fig. 4a). Masked 3D classification around the three-fold axis resulted in several classes that also appeared to further break the three-fold symmetry. Still, the quality of the obtained cryo-EM densities was poor, and we were unable to build atomic models or assign amino acid sequences to these extra internal densities (Supplementary Fig. 10). As expected, no CLD density was observed when performing the same analysis on the non-cargo-loaded *Sg* Enc shell (Supplementary Fig. 10), confirming that these densities represent part of the 2-MIBS CLD. The conserved hydrophobic residues F430, Y441, L285, and L264 are positioned below the observed CLD densities, and are nearly identical to the corresponding residues in the *A. baumannii* and *S. elongatus* Familiy 2A encapsulin three-fold CLD binding sites (Fig. 4b)^23,39^.

**Fig. 4.**
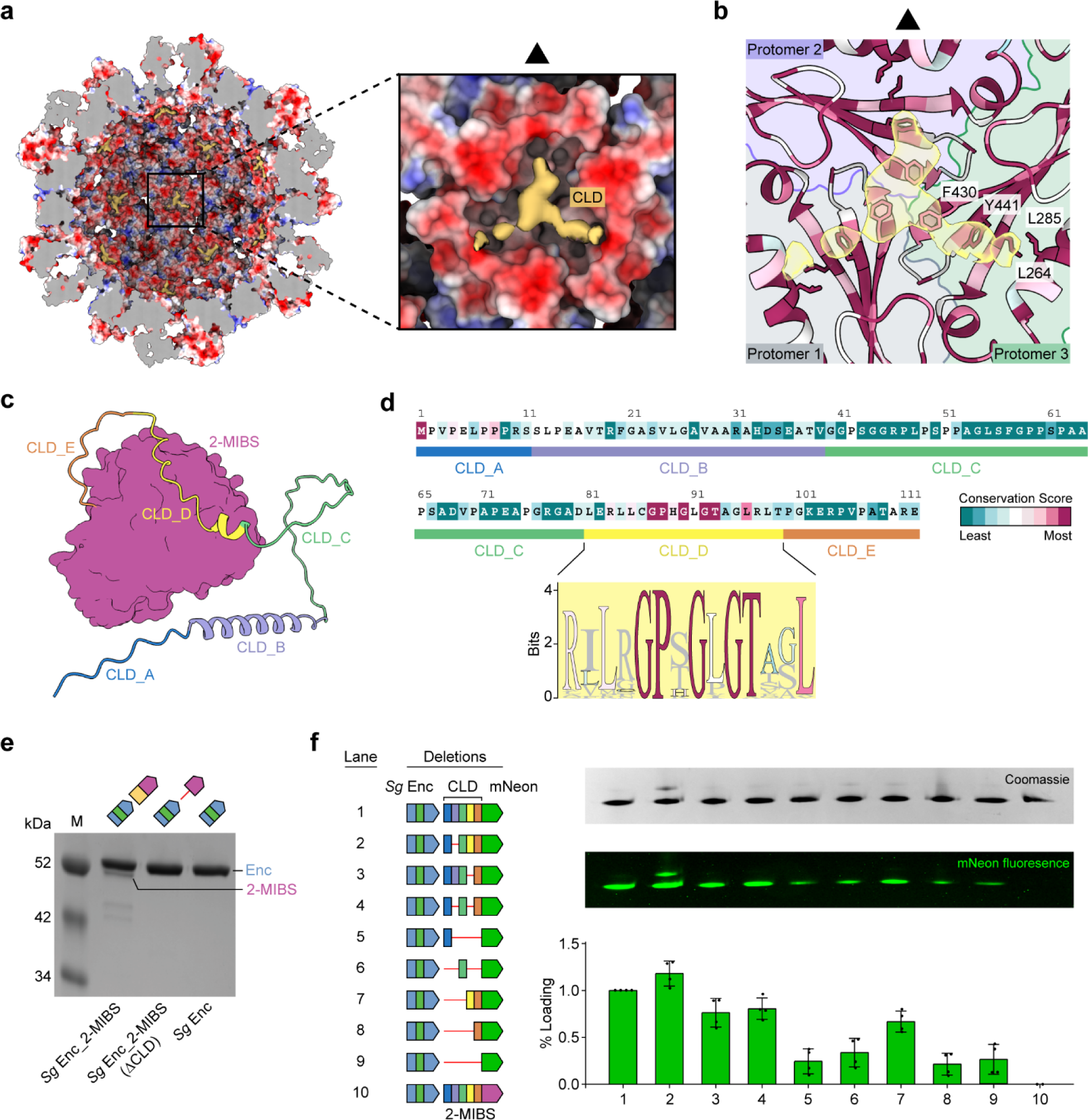
CLD-mediated cargo-loading of *Sg* Enc. **a**, Cut-way view of an *Sg* Enc shell model shown in electrostatic surface representation with extracted cryo-EM CLD densities obtained via asymmetric C1 refinement shown in yellow (left). Zoom-in highlighting CLD density bound to the interior of the shell at the three-fold axis of symmetry (right). **b**, The interior of *Sg* Enc surrounding the three-fold axis of symmetry shows strong sequence conservation throughout Family 2B encapsulins. Residues are colored based on ConSurf conservation analysis. The three protomers at the three-fold symmetry axis are highlighted by differently colored backgrounds. Conserved hydrophobic residues F430, Y441, L285, and L264 are shown as stick models and are located directly below the observed CLD density (transparent yellow). **c**, AlphaFold model of *Sg* 2-MIBS highlighting predicted secondary structure elements within the disordered N-terminal CLD. Five CLD elements – CLD_A to CLD_E – are shown in different colors. The structured TC domain of *Sg* 2-MIBS is shown in surface representation (purple). **d**, *Sg* 2-MIB CLD sequence colored by ConSurf conservation score. A sequence logo for the conserved motif within CLD_D is shown. **e**, SDS-PAGE analysis of purified *Sg* Enc shells. Deletion of the CLD abolishes cargo loading of *Sg* 2-MIBS when co-expressed with *Sg* Enc. *Sg* Enc_2-MIBS: 2-MIBS-loaded *Sg* Enc. ΔCLD: truncation mutant missing all of the CLD. **f**, Systematic deletion of the five identified CLD regions – CLD_A to CLD_E (left). A co-expression system of *Sg* Enc and CLD-fused mNeonGreen (mNeon), employed as a reporter, was used. Native PAGE analysis of purified truncation mutants shown on the right. Coomassie protein stain (right, top) and mNeonGreen fluorescence (right, middle) are shown. Normalized loading percentages based on gel densitometry and fluorescence analysis are shown (right, bottom). Data are shown as mean values, with error bars representing the standard deviation of four independent experiments.

The CLDs of Family 2 encapsulins are extended and flexible domains rich in proline, glycine, and alanine^22,23,39,43^. Inspection of *Sg* 2-MIBS AlphaFold models suggests two CLD stretches with helical propensity which we used as markers to subdivide the *Sg* 2-MIBS CLD into five regions – CLD_A to CLD_E – for mutational and cargo-loading analysis (Fig. 4c) ^64^. CLD_D contained the conserved consensus motif GPxGxGTxxL, mentioned above, which is the only conserved section of the *Sg* 2-MIBS CLD (Fig. 4d)^22,23^. Removal of the entire CLD from *Sg* 2-MIBS completely abolished cargo loading (Fig. 4e). To better understand the specific roles of the five CLD regions (CLD_A to CLD_E), we heterologously expressed a set of two-gene operons consisting of the *Sg* Enc shell protein and different variants of a CLD-mNeonGreen fusion protein containing different combinations of the five CLD regions (Fig. 4f). Analysis of purified *Sg* Enc_CLD-mNeonGreen variants via native PAGE allowed us to quantify cargo loading by comparing CLD-mNeonGreen fluorescence to the total protein content measured by Coomassie staining (Fig. 4f and Supplementary Fig. 11). Similar approaches have previously been successfully utilized to examine cargo loading in Family 1 and Family 2 encapsulin systems^23,65^. Removal of CLD_B resulted in slightly improved cargo-loading, while removal of all other CLD regions reduced cargo loading. Most notably, we observed that CLD_D, which contained the GPxGxGTxxL consensus motif, was not required for cargo loading, but did substantially improve cargo loading when present. Together with the observed, relatively low quality CLD cryo-EM densities, these results suggest that there are likely multiple redundant binding modes of CLD and its binding site on the interior of the shell. The CLD of *Sg* 2-MIBS is characterized by repetitive glycine-, alanine-, and proline-rich motifs that may contribute cooperatively to cargo-loading. This amino acid composition may enable multiple heterogeneous modes of binding that are not entirely sequence specific, but instead rely on physicochemical properties like flexibility and hydrophobicity for mediating CLD binding.

### *Sg* Enc is not a cAMP-binding protein

*Sg* Enc has previously been suggested to be involved in modulating antibiotic production in *Streptomyces coelicolor* A3(2) through binding of cAMP which is a well-known second messenger important for regulating secondary metabolism in *Streptomyces*^42^. As we now recognize *Sg* Enc to be a 2-MIBS-loaded encapsulin nanocompartment, we suspected that if *Sg* Enc is able to bind cAMP via its CBDs, this may result in a conformational change of the two-fold pores which in turn could control 2-MIB production by potentially restricting substrate flux across the encapsulin shell. To test this hypothesis, we monitored the activity of encapsulated and free 2-MIBS through a coupled assay with the GPP-methylating *Sg* MT in the presence or absence of cAMP^66-68^. We observed sigmoidal saturation curves (Supplementary Fig. 12), indicative of cooperative substrate binding, as well as substrate inhibition in conditions exceeding 90 μM GPP, features also observed for other terpene cyclases and prenyltransferases^66^. Apparent k_cat_/K_m_ values were consistent between free and encapsulated *Sg* 2-MIBS and the presence of 500 µM cAMP had no apparent influence on 2-MIBS activity (Supplementary Fig. 12). We suspect that the large two-fold pores of the *Sg* Enc shell allow rapid diffusion of substrates and products across the encapsulin shell causing encapsulation to have little influence on enzyme activity. GC-MS analysis further revealed that the presence of cAMP did not affect the product profile of *Sg* 2-MIBS, with 2-MIB as the only detectable product in our assays (Fig. 5b and Supplementary Fig. 13). 2-methylenebornane has been observed as a minor product of 2-MIBS in *in vitro* assays^17^, yet we did not detect it in any of our assays. These results show that cAMP has no effect on the activity or product profile of encapsulated *Sg* 2-MIBS.

**Fig. 5.**
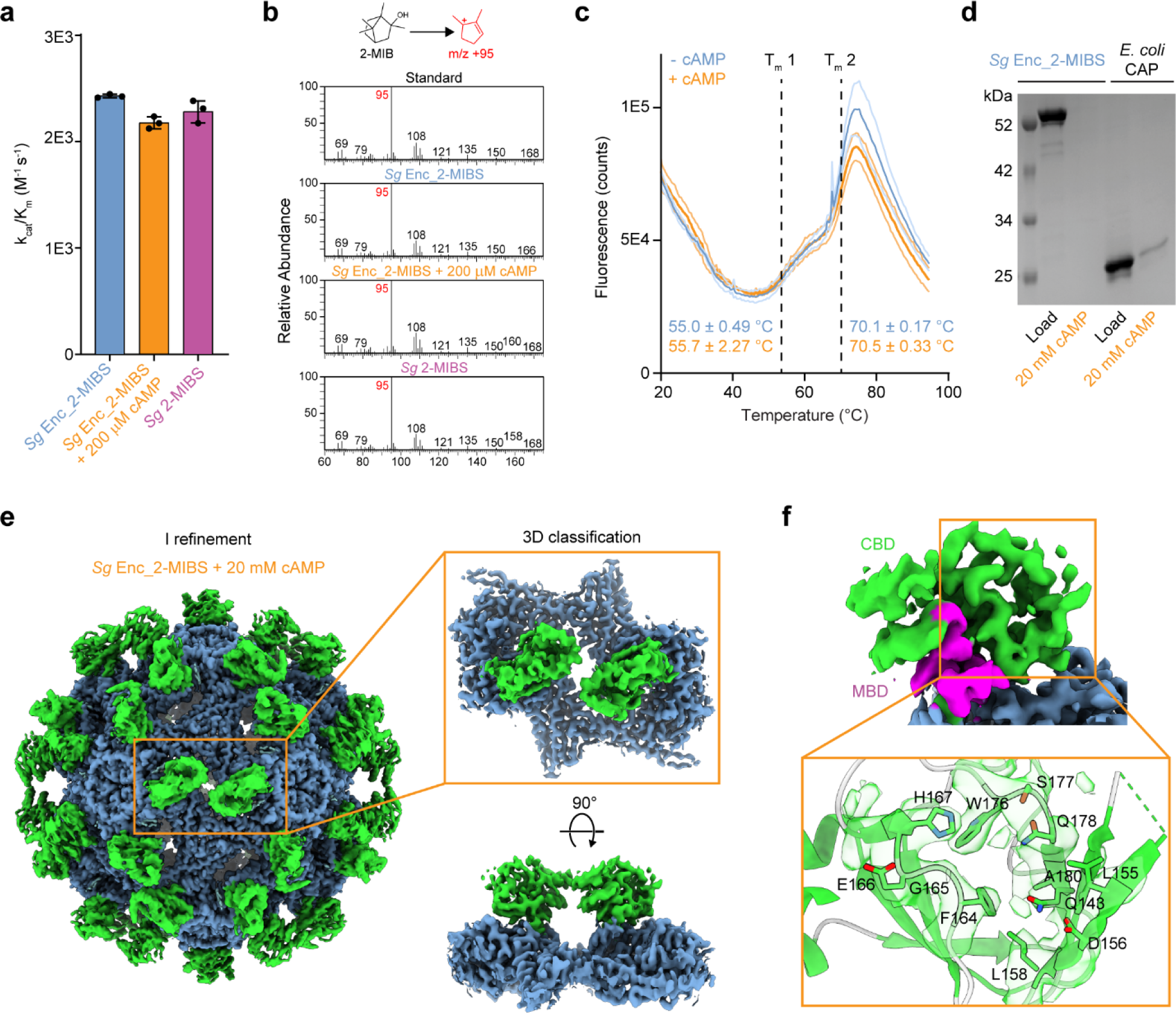
The effect of cAMP on *Sg* 2-MIBS activity and cAMP binding to *Sg* Enc CBD. **a,** *In vitro* activity assays of 2-MIBS-loaded *Sg* Enc in the absence and presence of cAMP. Free 2-MIBS activity is shown for reference. k_cat_/K_m_: catalytic efficiency of 2-MIBS. Data are shown as mean values, with error bars representing the standard deviation of three independent experiments. **b,** GC-MS analysis of 2-MIBS activity assays. All assays yielded 2-MIB as the main product, with its diagnostic mass fragment (m/z +95) identified as the most abundant ion. A 2-MIB standard is shown for reference (top). **c,** Differential scanning fluorimetry (DSF) thermal melts of 2-MIBS-loaded Sg Enc in the absence or presence of cAMP. T_m_: melting temperature. **d,** 2-MIBS-loaded Sg Enc binding to cAMP-modified agarose resin. A representative SDS-PAGE gel is shown demonstrating that 2-MIBS-loaded *Sg* Enc is unable to bind to cAMP-modified agarose resin. The positive control *Ec* CAP is able to bind to the resin. Load: sample loaded onto the resin. 20 mM cAMP (orange): elution from the resin with 20 mM cAMP. **e,** Cryo-EM analysis of *Sg* Enc in the presence of cAMP. No significant conformational changes of the CBDs or the HK97-domain could be observed when compared to cryo-EM maps obtained in the absence of cAMP. Improved CBD density was obtained via 3D classification followed C1 homogeneous refinement, but no conformational changes could be observed. **f**, Cryo-EM density of the β-barrel CBD binding site is shown (top). No additional density could be observed in the presence of cAMP. The metal-binding subdomain (MBD) is shown in purple. Structural model of the binding site with binding site residues shown as stick (bottom).

We next sought to directly investigate cAMP binding to *Sg* Enc CBDs. Differential scanning fluorimetry (DSF), a method commonly used to measure ligand binding^69^, was initially carried out and identified two thermal transitions for cargo-loaded *Sg* Enc shells at melting temperatures (T_m_) of 55.0 ± 0.49°C and 70.1 ± 0.17°C (Fig. 5c). Addition of cAMP yielded nearly identical thermal transitions, with T_m_ values of 55.7 ± 2.27°C and 70.5 ± 0.33°C, highlighting that cAMP addition does not result in any observable stabilization of the *Sg* Enc shell. Using an orthogonal cAMP binding assay, we next employed a cAMP-modified agarose resin, another common strategy for purifying and identifying cAMP-binding proteins, to test cargo-loaded *Sg* Enc cAMP binding capacity^70-73^. No binding to cAMP-agarose resin could be observed for *Sg* Enc shells while *Ec* CAP – a known cAMP-binding protein – exhibited clear resin binding (Fig. 5d). Subsequently, cryo-EM analysis of 2-MIBS-loaded *Sg* Enc was carried out in the presence of an excess of 20 mM cAMP to potentially capture a cAMP-bound state (Supplementary Fig. 14). Yet, we observed no apparent conformational changes in cryo-EM maps resulting from I refinements when compared to maps obtained in the absence of cAMP (Fig. 5e). Masked 3D classification and homogeneous C1 refinements of CBDs resulted in cryo-EM maps with improved CBD density but showed no significant conformational changes compared to previous maps (Fig. 5f). Closer analysis of the CBD revealed an empty binding pocket, identical to the binding pockets observed without any cAMP present. In sum, these orthogonal analyses demonstrate that cAMP is likely not a native ligand of *Sg* Enc CBDs and that addition of cAMP has no observable influence on shell structure or CBD conformation.

## Discussion

In this study, we demonstrate that *Sg* Enc, formerly known as EshA, is not a transcription factor or cAMP-binding protein as previously suggested^20,40-42^, but is instead a Family 2B encapsulin shell protein that functions to sequester the terpene cyclase 2-MIBS. Our characterization of *Sg* Enc experimentally confirms that encapsulin nanocompartments are directly involved in secondary metabolite biosynthesis and that the ubiquiteous odorant 2-MIB is biosynthesized inside a protein shell in many soil bacteria. Family 2B encapsulins and their respective gene clusters are widespread among Actinobacteria, Proteobacteria, and Cyanobacteria and, based on computational searches, have been hypothesized to encapsulate a variety of enzymes associated with secondary metabolism, including diverse terpene cyclases, polyprenyl transferases, and xylose-modifying enzymes^22,23^. Additionally, many Family 2B encapsulin-encoding gene clusters encode multiple putative cargo proteins, potentially resulting in protein shells loaded with multiple types of cargo^22^. Further, some Family 2B encapsulin operons seem to encode two distinct shell protein copies, suggesting that these systems may be able to assemble into two-component encapsulin shells^22^.

Our structural analysis of the 2-MIBS-loaded *Sg* Enc shell reveals that the *Sg* Enc shell protein exhibits an HK97-like fold similar to Family 2A encapsulins^23,39^, but contains large and elongated two-fold pores not seen in any other encapsulin shell. These two-fold pores in the *Sg* Enc shell are potentially necessary for substrate and product flux across the shell given the relatively large size of the 2-MIBS substrate – 2-methylGPP – and product – 2-MIB. However, large two-fold pores may not be a feature of all Family 2B encapsulin shells as approximately 9% of Family 2B systems are associated with cysteine desulfurase cargos that utilize the small substrate L-cysteine^22^. In characterized Family 2A encapsulins, lacking CBDs, that encapsulate cysteine desulfurases, the two-fold pores are essentially closed by the structured N-arms of two neighboring shell protein protomers^23,39^. N-arms of Family 2B encapsulins display high sequence and length variability, which may allow different alternate orientations of the N-arms and consequently different two-fold pore states which may be static or dynamic^22^. Dynamic five-fold pores have recently been reported in Family 1 encapsulins^74-76^. Furthermore, as mentioned above, some Family 2B gene clusters encode for two distinct Family 2B encapsulin shell proteins, which may form heterogeneous two-component shells with a mixture of open and closed two-fold pores, depending on the respective N-arm states of the constituent shell protomers.

Our analysis of the *Sg* 2-MIBS CLD demonstrates its importance for cargo loading, as predicted by previous bioinformatic analyses and related experimental evidence from Family 2A encapsulin systems^22,23,39^. Despite considerable sequence variability, 2-MIBS CLDs are rich in glycine, alanine, and proline, a feature shared with Family 2A CLDs^22,23,39^. Mutational analysis of the *Sg* 2-MIBS CLD suggests that repetitive stretches of glycine, alanine, and proline may provide redundant interaction motifs that mediate CLD-shell binding. Our cryo-EM analysis revealed interactions between the *Sg* 2-MIBS CLD and the interior surface surrounding the three-fold axis of the *Sg* Enc shell. The observed variability and limited resolution of these CLD densities may be a result of multiple heterogenous modes of binding, resulting from redundant CLD interaction motifs^39^. Given the high sequence and length variability observed throughout 2-MIBS CLDs, other 2-MIBSs with longer CLDs may utilize additional binding interfaces extending beyond the three-fold axis. Inversely, 2-MIBSs found in branch 2 of our phylogenetic tree (Fig. 1c) may contain minimal binding motifs with more homogeneous three-fold binding modes.

The most unique feature of Family 2B encapsulins is the presence of an insertion domain within the encapsulin shell protein, annotated as a CBD. Our structural analysis of the *Sg* Enc CBD reveals that it adopts a fold highly similar to other characterized cAMP-binding CBDs^44^. In addition, we identified a unique and conserved metal-binding subdomain (MBD) as part of the CBD which likely contributes to stabilizing CBD orientation through an extensive interface with the HK97-like domain of the Family 2B encapsulin shell protomer. We were unable to observe any conformational changes of the CBDs or HK97-like domain of *Sg* Enc, regardless of cargo-loading state or the presence of cAMP. Diverse ligand binding and cryo-EM experiments suggest that *Sg* Enc CBDs do not bind cAMP. We hypothesize that this may be partially caused by the presence of a conserved tryptophan (W176) – replacing a conserved arginine found in true cAMP-binding proteins – in the putative binding pocket, a feature widespread among Family 2B encapsulin CBDs. While the conserved arginine of confirmed cAMP-binding proteins is crucial for binding the cyclic phosphate of cAMP, the tryptophan residue found in Family 2B CBDs would not be able to mediate a similar binding interaction with cAMP^22^. A number of examples exist of cAMP-insensitive CBDs, able to bind other ligands^54,56-59,77-79^. Based on our results, we hypothesize that the CBDs of the *Sg* Enc shell may be able to bind to an as yet unidentified, non-cAMP ligand. If the *Sg* Enc CBD behaves like many other CBDs, ligand binding may trigger a conformational change in the *Sg* Enc shell which could function to regulate the activity of internalized cargo in a ligand-dependent fashion.

Our results redefine the function of EshA and experimentally establish Family 2B encapsulins as cargo-loaded protein nanocompartments involved in secondary metabolite biosynthesis. A number of key questions remain: What is the molecular logic and advantage of 2-MIBS encapsulation? What is the native ligand of *Sg* Enc CBD and how are CBDs involved in mediating or regulating 2-MIB biosynthesis? Our findings provide a solid foundation for future investigations into these questions, and also provide molecular level detail to aid in adapting Family 2B encapsulin nanocompartments for future synthetic biology and bionanotechnology applications.

## Methods

### Computational and phylogenetic analyses

An initial sequence similarity network (SSN) for 2-MIBS was created using the EFI-EST server^80,81^ in January of 2023. The amino acid sequence of 2-MIBS from *S. griseus* (Uniprot ID: A0A2X2LWM5) was used as a BLAST query in EFI-EST with an E-value of -5, a maximum of 5,000 sequence hits, and minimum sequence length of 300 amino acids. This initial step returned 990 putative 2-MIBS encoding genes whose genome neighborhoods were then visualized using the EFI-GNT server^80,81^. To distinguish 2-MIBS-encoding genes from other similar single domain terpene cyclases, only genome neighborhoods containing MT-encoding genes (Pfam IDs: PF02353, PF08241, PF13649) within 10 kb of the putative 2-MIBS genes were included in the dataset, resulting in 745 unique 2-MIBS containing gene clusters. The dataset was then manually curated to remove spurious entries, resulting in a final dataset of 734 unique 2-MIBS-encoding gene clusters. 723 gene clusters identified from Actinobacteria were then used to assemble a phylogenetic tree, using the NGphylogeny.fr server,^82^ via MAFFT alignment^83^ followed by phylogenetic tree assembly using the FastTree algorithm^84,85^ with 1,000 bootstraps^86^. The tree was then visualized and annotated using iTOL v6^87^.

For SSN analysis of the 2-MIBS proteins contained within the final dataset, the EFI-EST server^80,81^ was used to assemble an initial SSN, which was then annotated and modified using Cytoscape v3.1.0^88^ to keep only edges with at least 65% sequence identity. Sequence alignments of the CLDs were performed using Clustal Omega on the EMBL-EBI server^89^. Sequence logos were visualized using Geneious Prime version 2023.0.4.

### Molecular cloning

Plasmids for protein expression in *S. coelicolor* A3(2) were assembled according to previously published methods^90^. Briefly, to construct the *Sg* Enc_2-MIBS expression plasmid, genes encoding for the *Sg* Enc (mmpI_1) and *Sg* 2-MIBS (NCTC13033_02227) were amplified as a single amplicon from *S. griseus* subs. *griseus* (ATCC 23345) genomic DNA and adding NdeI and BamHI overhangs. The amplicon and a pGM1190 vector were cut using NdeI (ThermoScientific, Cat # FD0583) and BamHI (ThermoScientific, Cat # ER0051) restriction enzymes, purified by extraction from an agarose gel. The plasmid was subsequently assembled using T4 ligase (New England Biosciences, Cat # M0202S). *E. coli* BL21 (DE3) cells were then transformed with the assembled plasmid via electroporation and grown on LB agar plates containing 50 µg/mL of apramycin at 37°C for 18 h. Plasmid-containing colonies were screened by Sanger sequencing (Eurofins Scientific). The confirmed plasmid was then used as a template for cloning *Sg* Enc*, Sg* Enc_ΔCLD-2-MIBS, and *Sg* Enc_CLD-mNeonGreen constructs, which were cloned by either Gibson assembly or restriction cloning. For the *Sg* Enc, *Sg* 2-MIBS, *Sg* MT, and *Ec* CAP constructs expressed in *E. coli* BL21 (DE3), codon-optimized gBlocks from IDT were used and designed to contain 20 base pair overhangs corresponding to the NdeI and PacI cut sites of the pETDuet-1 vector. The *Sg* MT and *Ec* CAP gBlocks were designed to encode an 8xHi-tag followed by a tobacco etch protease (TEV) cleavage site at the N-terminus, whereas the *Sg* 2-MIBS plasmid was designed to encode a C-terminal TEV site followed by an 8xHis-tag. The pETDuet-1 vector was linearized using NdeI and PacI (ThermoScientific, Cat # FD2204), and purified by extraction from an agarose gel. The plasmids were then assembled via Gibson assembly (New England Biolabs, Cat # E2611L), followed by transformation of *E. coli* BL21 (DE3) cells via electroporation. Cells were plated on LB agar plates containing 100 µg/mL of ampicillin and grown at 37°C for 18 h. Plasmids were then confirmed by Sangar sequencing.

### Protein expression

*Sg* Enc, *Sg* 2-MIBS, *Sg* MT, and *Ec* CAP were expressed in *E. coli* BL21 (DE3) cells by inoculating 0.5 to 1 L of LB media containing 100 μg/mL of ampicillin with an overnight starter culture, followed by shaking at 200 rpm and 37°C until the cells reached an OD_600_ of 0.5. Protein expression was then induced by the addition of 0.1 mM IPTG for *Sg* Enc and 0.5 mM IPTG for *Sg* 2-MIBS, *Sg* MT, and *Ec* CAP, followed by shaking at 200 rpm and 18°C for 18 h. Cells were harvested, and cell pellets were stored at -20°C until use.

Heterologous protein expression in *S. coelicolor* A3(2) (ATCC #BAA-471, also known as *Streptomyces violaceoruber* strain John Innes Centre M145) was performed according to previously published protocols^90,91^. *E. coli* ET12567/pUZ8002 cells were transformed with pGM1190 vectors containing the *Sg* Enc_2-MIBS, *Sg* Enc, *Sg* Enc_ΔCLD-2-MIBS, and *Sg* Enc CLD-mNeonGreen constructs via heat shock and maintained on LB agar plates containing 50 μg/mL apramycin, 30 μg/mL chloramphenicol, and 50 μg/mL kanamycin. Plasmid-containing colonies were then used to inoculate 20 mL overnight cultures of LB containing 50 μg/mL apramycin, 30 μg/mL chloramphenicol, and 50 μg/mL kanamycin, which were grown overnight at 37°C with shaking at 200 rpm. The overnight cultures of *E. coli* ET12567/pUZ8002 were harvested the following morning by centrifugation at 3,800 x g for 10 min. Cells were washed by resuspending the cell pellets in 20 mL LB followed by centrifugation at 3,800 x g for 10 min. This step was repeated again, and the final cell pellet was then resuspended in 2 mL of LB. 200 μL of plasmid-containing resuspended *E. coli* ET12567/pUZ8002 cells were then mixed with 500 µL of freshly prepared *S. coelicolor* A3(2) spore stock in GYM medium which had been heated at 50°C for 10 min immediately prior to mixing with *E. coli* ET12567/pUZ8002 cells. The cell mixtures were then plated on MS agar containing 10 mM MgCl_2_, and incubated overnight at 30°C. The following morning, the plates were overlaid with 1 mg nalidixic acid and 1 mg of apramycin and incubated at 30°C for 3 to 5 days until spores became visible. Individual colonies with visible spores were then either struck or plated as lawns on fresh plates of MS agar containing 50 μg/mL apramycin and 25 μg/mL nalidixic acid. The plates were then incubated for another 3 to 5 days and 30°C until spores were visible. The spores were scraped from the plates and used to inoculate 500 mL baffled flasks containing YEME media with glass beads and 50 μg/mL apramycin. The cultures were grown by shaking at 200 rpm at 30°C until the OD_600_ reached 0.4 to 0.8, which typically took 24 to 36 h. To induce protein expression, 20 μg/mL of thiostrepton was added to the media, and cells were grown with shaking at 200 rpm and 30°C for 72 h. Cells were then harvested by centrifugation and stored at -20°C until further use.

### Protein purification

*Sg* Enc_2-MIBS for cryo-EM analysis was purified according to previously published protocols^90^. 1.5 g of a frozen *S. coelicolor* A3(2) cell pellet was resuspended in 7.5 mL of lysis buffer consisting of 150 mM NaCl, 20 mM Tris pH 7.5, 2 mM MgCl_2_, 1 mg/mL lysozyme, 250 units of Benzonase nuclease (Millipore Sigma), and a protease cocktail consisting of 10 µM leupeptin, 14 µM E-64, 0.4 µM aprotinin, 0.4 mM AEBSF, and 114 µM bestatin. The resuspended cell pellet was then lysed by sonication for 3 min on ice, pulsing for 10 s on and 20 s off, with a power of 96 W using a Model 120 Sonic Dismembrator (Fisher Scientific, Inc). The lysate was then centrifuged at 21,000 x g for 10 min to remove cell debris. Ammonium sulfate was added to the clarified supernatant to 50% saturation, and the supernatant was rocked at 4°C for 1 h. Precipitated protein was pelleted by centrifugation at 10,000 x g and 4°C for 10 min. The protein pellet was resuspended in 10 mL of 150 mM NaCl, 2 mM MgCl_2_, 20 mM Tris pH 7.5, and concentrated to 0.5 mL using an Amicon 100 Ultra-15 centrifugal filter (Millipore) with a molecular weight cutoff of 100 kDa. Approximately 125 units of Benzonase nuclease were added to the concentrated protein and the mixture incubated at 4°C for 30 min. The protein sample was then loaded onto an AKTA Pure FPLC system and purified with a Superose 6 Increase 10/300 GL column using 150 mM NaCl, 20 mM Tris pH 7.5 as the running buffer. *Sg* Enc_2-MIBS containing fractions were pooled, concentrated to approximately 3 mg/mL, and stored at 4°C until use.

*E. coli*-expressed *Sg* Enc for cryo-EM analysis was purified by resuspending approximately 3 g of cell pellet in 15 mL of lysis buffer. The resuspended pellet was then lysed by sonication for 3 minutes on ice, pulsing for 10 s on and 20 s off, at a power of 75 W. The lysate was clarified by centrifugation at 8,000 x g and 4°C for 10 min. 10% (w/v) PEG 8,000 and 450 mM NaCl were added to the clarified lysate, which was further rocked at 4°C for 1 h to precipitate *Sg* Enc shells. The lysate was then clarified by centrifugation at 8,000 x g and 4°C for 10 min. The supernatant was discarded, and the pellet was resuspended in a minimal volume of 150 mM NaCl, 20 mM Tris pH 7.5 and filtered using a 0.22 µm syringe filter. The clarified protein sample was then loaded onto an AKTA Pure FPLC and purified with a Sephacryl S-500 size exclusion column. *Sg* Enc-containing fractions were then pooled, and subsequently buffer exchanged into 20 mM Tris pH 7.5 buffer using an Amicon Ultra-15 100 kDa centrifugal filter, loaded onto an AKTA Pure FPLC system, followed by purification with a HiPrep DEAE FF 16/10 column using a linear gradient from 0 to 1 M NaCl, 20 mM Tris pH 7.5. This step was performed twice to better enrich the *Sg* Enc protein sample. *Sg* Enc-containing fractions were pooled, concentrated to 0.5 mL, and loaded onto an AKTA Pure FPLC and purified using a Superose 6 10/300 GL column with 150 mM NaCl, 20 mM Tris pH 7.5 as the running buffer. *Sg* Enc containing fractions were then pooled, concentrated to 3.0 mg/mL, and stored at 4°C until further use.

*S. coelicolor* A3(2)-expressed *Sg* Enc for cryo-EM analysis was purified using the same method as for *S. coelicolor* A3(2)-expressed *Sg* Enc_2-MIBS outlined above, however, the *Sg* Enc sample was further purified through an additional step by loading onto an AKTA Pure FPLC system and purifying via a HiTrap Q-FF column using a gradient from 20 mM Tris pH 8.0, 5 mM MgCl_2_ to 20 mM Tris pH 8.0, 5 mM MgCl_2_, 1 M NH_4_Cl. *Sg* Enc-containing fractions were then pooled, concentrated to 3.2 mg/mL, desalted into 150 mM NaCl, 20 mM Tris pH 7.5, and stored at 4°C until use.

*S. coelicolor* A3(2)-expressed *Sg* Enc_2-MIBS for cryo-EM analysis in the presence of 20 mM cAMP was purified by resuspending 3 g of frozen *S. coelicolor* A3(2) cell pellet in a lysis buffer consisting of 20 mM tris pH 8.0, 10 mM MgCl_2_, 20 mM ammonium acetate, 1x protease inhibitor cocktail as above, 250 U of Benzonase nuclease, and 1 mg/mL of lysozyme. The resuspended cell pellet was then lysed by sonication for 3 min on ice, pulsing for 10 s on and 20 s off, with a power of 96 W. The lysate was then centrifuged at 21,000 x g for 10 minutes to remove cell debris. Ammonium sulfate was then added to the clarified supernatant to 50% saturation, and the supernatant was rocked at 4°C for 1 h. Precipitated protein was then pelleted by centrifugation at 10,000 x g for 10 min at 4°C. The protein pellet was resuspended in 10 mL of 20 mM Tris pH 8.0, 10 mM MgCl_2_, 20 mM ammonium acetate, and concentrated to 0.5 mL using an Amicon 100 Ultra-15 100 kDa filter. Approximately 125 units of Benzonase nuclease were added to the concentrated protein followed by incubation at 4°C for approximately 30 min. The protein sample was then loaded onto an AKTA Pure FPLC and purified using a Superose 6 Increase 10/300 GL column in 20 mM Tris pH 8.0, 10 mM MgCl_2_, 20 mM ammonium acetate running buffer. *Sg* Enc_2-MIBS-containing fractions were pooled, again loaded onto an AKTA Pure FPLC, and purified via a HiTrap Q-FF column using a gradient from 20 mM Tris pH 8.0, 5 mM MgCl_2_ to 20 mM Tris pH 8.0, 5 mM MgCl_2_, 1 M NH_4_Cl. *Sg* Enc_2-MIBS-containing fractions were pooled, desalted into 20 mM Hepes pH 7.5, 5 mM MgCl_2_ using an Amicon 100 Ultra-15 100kDa filter, loaded onto an AKTA Pure FPLC, and purified via a HiTrap S-FF column using a gradient from 20 mM Hepes pH 7.5, 5 mM MgCl_2_ to 20 mM Hepes pH 7.5, 5 mM MgCl_2,_ 1M NH_4_Cl. *Sg* Enc-containing fractions were then pooled, concentrated to 0.5 mL using an Amicon 100 Ultra-15 100 kDa filter, again loaded onto an AKTA Pure FPLC, and purified using a Superose 6 Increase 10/300 GL column equilibrated with 20 mM Hepes pH 7.5, 25 mM NaCl, 5 mM MgCl_2_. *Sg* Enc_2-MIBS-containing fractions were pooled, concentrated, and exchanged into 20 mM Hepes pH 7.5, 5 mM MgCl_2_. cAMP was added to a final concentration of 20 mM approximately 1 h before cryo-EM grids were prepared with a final protein concentration of 3.5 mg/mL. During this purification, the cation and anion exchange purification steps were performed to remove any non-specifically bound charged ligands.

*S. coelicolor* A3(2)-expressed *Sg* Enc_2-MIBS for enzymatic assays and cAMP binding experiments was purified by the same method used to prepare *S. coelicolor* A3(2)-expressed *Sg* Enc_2-MIBS for cryo-EM analysis in the presence of cAMP, except for omitting the Hi-Trap S-FF purification step. Similarly, *S. coelicolor* A3(2)-expressed *Sg* Enc_CLD-mNeonGreen variants were purified using the same methods, except for omitting both the Hi-Trap S-FF and second Superose 6 Increase 10/300 GL purification steps. For enzymatic assays, *Sg* Enc_2-MIBS was stored at 4°C until use. *Sg* Enc_2-MIBS used for product profile experiments, DSF, and cAMP-agarose binding experiments was drop frozen in liquid nitrogen and stored at -80°C until use. *Sg* Enc_CLD-mNeonGreen variants were also drop frozen in liquid nitrogen and stored at - 80°C until use.

*E. coli*-expressed *Sg* 2-MIBS was purified by resuspending 4 g of frozen cell pellet in 20 mL of 20% glycerol, 300 mM NaCl, 20 mM imidazole pH 8.0, 20 mM Tris pH 8.0, 5 mM MgCl_2_, 1 x protease inhibitor cocktail used in all other purifications, and 0.5 mg/mL lysozyme, followed by sonication on ice at 88 W for 3 min, in pulses of 10 s on separated by 20 s off. The lysate was centrifuged for 10 min at 21,000 x g and the clarified supernatant loaded onto an AKTA Pure FPLC system and purified via a His-Trap FF column using a gradient from 20% glycerol, 300 mM NaCl, 20 mM imidazole pH 8.0, 20 mM Tris pH 8.0, 5 mM MgCl_2_ to the same buffer but containing 1M imidazole. *Sg* 2-MIBS-containing fractions were pooled, concentrated to 0.5 mL in an Amicon Ultra-15 centrifugal filter with a 30 kDa molecular weight cutoff, again loaded on an AKTA pure FPLC, and purified via a Superdex S200 column in 10 mM Hepes pH 7.5, 5 mM MgCl_2_ running buffer. *Sg* 2-MIBS-containing fractions were pooled, concentrated to approximately 5 mg/mL and stored at 4°C if used for enzymatic activity assays, otherwise, remaining protein was drop frozen in liquid nitrogen and stored at -80°C.

*E. coli*-expressed *Sg* 2-MT was purified using the same lysis buffers and initial His-Trap FF step as used for the *Sg* 2-MIBS purification. Following this initial step, 3 mg of recombinantly-expressed tobacco-etch virus (TEV) protease was added to *Sg* MT, which was then dialyzed against 1 L of 20 mM Tris pH 8.0, 300 mM NaCl, 20% glycerol overnight at 4°C to remove the cleaved N-terminal 8xHis-tag. Then, the protein sample was again purified via a HisTrap-FF column, with TEV-cleaved protein collected in the flow-through, exchanged into 20 mM Tris pH 8.0, 300 mM NaCl, 20% glycerol, 1 mM DTT using an Amicon Ultra-15 100 kDa filter unit, and concentrated to 20 mg/mL. The protein was then drop frozen in liquid nitrogen and stored at -80°C until further use.

*E. coli*-expressed *Ec* CAP was purified using the same lysis buffers as for *Sg* MT and *Sg* 2-MIBS and initial His-Trap FF step. Following the His-Trap-FF step, *Ec* CAP was dialyzed overnight with stirring against 1 L of 50 mM Na_2_/KPO_4_ pH 7.0, 300 mM NaCl, and 0.1 mM TCEP at 4°C. *Ec* CAP was then concentrated to ca. 2.5 mg/mL using an Amicon Ultra-15 centrifugal filter unit with a molecular weight cutoff of 10 kDa, drop frozen in liquid nitrogen, and stored at -80°C until use.

### Negative stain transmission electron microscopy (TEM)

Protein samples were diluted to 0.1 to 0.15 mg/mL and applied to freshly glow discharged (60 s, 5 mA, PELCO easiGlow) 200 mesh gold formvar/carbon square mesh grids (EMS, Cat # FCF200-AU-EC). After 60 s, the samples were blotted of the grids using filter paper. Grids were then washed with a 5 µL drop of distilled water, followed by two rounds of 5 s and then 30 s staining with 5 µL drops of 0.75% uranyl formate. TEM micrographs were recoreded using an FEI Morgagni TEM operating at 100 kV located at the Michigan University Life Sciences Institute.

### Native PAGE analysis

Approximately 1 µg of purified *Sg* Enc-CLD-mNeonGreen variants, were loaded onto Invitrogen Native PAGE 3-12% Bis-Tris gels. The native PAGE gels were run using 1x Invitrogen Native PAGE buffer for 120 min at 150 V at 4°C. In-gel mNeonGreen fluorescence was imaged using a GelDoc Go imaging system (BioRad Laboratories, Inc) equipped with a blue sample tray. Gels were then stained overnight using Ready Blue Stain (Sigma Aldrich), destained briefly in water, and imaged using a ChemiDoc Imaging system (BioRad Laboratories, Inc). Protein bands on Coomassie-stained and fluorescently imaged Native PAGE gels were analyzed via gel densitometry in ImageJ2^92^. Protein amount per lane was normalized to the *Sg* Enc-CLD-mNeonGreen sample. The same approach was taken to obtain a normalized mNeonGreen fluorescence intensities for each sample. Relative cargo-loading was then calculated as a ratio of normalized protein amount and normalized fluorescence intensiy.

### 2-MIBS activity assays

2-MIBS activity assays were performed by pre-incubating samples of 10 to 200 µM GPP (Sigma Aldrich, Cat # 76532-10MG) in a buffer containing 25 µM purified *Sg* MT, 2 mM SAM (Sigma Aldrich, Cat # A7007-25MG), 10 mM MgCl_2_, and 20 mM Hepes pH 7.5 for 2 h at 30°C to convert GPP to 2-methyl-GPP. Next, 2-amino-6-mercapto-7-methyl purine ribonucleoside (MESG), purine nucleoside phosphorylase (PNP), and inorganic pyrophosphatase (PPase) were added according to the EnzCheck pyrophosphate assay kit (ThermoFisher, Cat # E-6645). The samples were briefly centrifuged for 3 min at 10,000 x g to remove any precipitated *Sg* MT, and then 30 µL of each sample was aliquoted in triplicate into 384 well plates (Corning, Cat # 3540). Assays were initiated by addition of 10 µL of 2 µM purified free *Sg* 2-MIBS, or 10 µL of 20 µM purified encapsulated 2-MIBS (*Sg* Enc_2-MIBS) to each sample well, resulting in final assay volume of 40 µL and final concentrations of 5 to 100 µM GPP/2-methyl-GPP, 6.25 µM *Sg* MT, 0.5 mM SAM/SAH, 5 mM MgCl_2_, 10 mM Hepes pH 7.5, 200 µM MESG, 0.04 U PNP, and 0.0012 U PPase. Absorbance at 360 nm, corresponding to the conversion of MESG to ribose 1-phosphate and 2-amino-6-mercapto-7-methylpurine by PNP, were collected every 22 s in sweep mode using a BioTek Synergy H1 microplate reader. Slopes were calculated in the linear reaction periods (8 to 14 min) and used to calculate reaction velocities using a pyrophosphate standard curve. Velocities were plotted and curves were fitted to an allosteric sigmoidal model in GraphPad Prism 9.10.

### Gas chromatography mass spectrometry (GC-MS) analysis

Reactions were prepared in sealed vials containing 1 mM SAM, 75 µM GPP, 12.5 µM *Sg* MT, 10 mM Hepes pH 7.5, 5 mM MgCl_2_, 1 mM DTT and 0.5 µM *Sg* 2-MIBS, 5 µM *Sg* Enc_2-MIBS, or 5 µM *Sg* Enc_2-MIBS + 500 µM cAMP. A 100 µL pentane cushion was pipetted on the top of each reaction, and the reactions were incubated for 20 h at room temperature. Samples were centrifuged for 1 min at 14,000 x g, the aqueous layer was transferred to a tube containing 200 µL of pentane, mixed by vortexing, and spun again for 1 min at 14,000 x g, and the pentane layer then pooled with the other pentane layers. 2 to 3 small crystals of disodium sulfate were added to each pentane extract to remove residual water, incubated for 10 min at room temperature, and the pentane layer then transferred to sample vials for GC-MS analysis. The same extraction procedure was used for the 2-MIB standard with a starting concentration of approximately 100 µM. 1 µL of each sample was injected into a Shimadzu QP-2010 GC-MS instrument equipped with a DB-5 column, run in splitless mode with a column flow rate of 0.97 mL/min and a temperature ramp of 60°C to 280°C at 20°C/min. Data analysis was carried out in OpenChrom v 1.5.

### Differential scanning fluorimetry (DSF)

Thermal melts of *Sg* Enc_2-MIBS or *Sg* Enc_2-MIBS + 200 µM cAMP were obtained by DSF using an Uncle instrument (Unchained Labs). DSF samples contained 10x SYPRO orange dye (diluted from 5000x stock in DMSO, Invitrogen, Cat # S6651), 1 µM *Sg* Enc_2-MIBS, 10 mM Hepes pH 7.5, 0.1 mM TCEP, with or without additional 200 µM cAMP. Thermal ramps progressed from 20°C to 95°C at a rate of 1°C/min, and were monitored by measuring fluorescence emission spectra from 250 to 720 nm with an excitation wavelength of 473 nm. The Uncle Analysis V5.04 software package was used to calculate the area under the curve between 510 and 680 nm, which was used to represent thermal transition curves and calculate differential inflection points representative of thermal transitions (melting temperature, T_m_).

### cAMP-agarose binding assays

cAMP-agarose binding assays were carried out in batch format. 0.2 mg of purified *Sg* Enc_2-MIBS or *Ec* CAP was exchanged into binding buffer consisting of 100 mM NaCl, 10 mM Hepes pH 7.5, and 5 mM MgCl_2_. 300 µL of *Sg* Enc_2-MIBS or *Ec* CAP was then incubated with 100 µL of cAMP-Agarose (Sigma Aldrich, Cat # A0144) in the same buffer for 15 min at room temperature with gentle rocking. The resin was centrifuged for 3 min at 3,000 x g, the supernatant was discarded, and the resin was resuspended in 500 µL of binding buffer followed by centrifugation at 3,000 x g. The supernatant was discarded, and the wash step was repeated two more times to remove non-specifically bound protein. Bound protein was eluted by resuspending the resin twice in 200 µL of 20 mM cAMP, 100 mM NaCl, 10 mM Hepes pH 7.5, 5 mM MgCl_2_.

### Cryo-electron microscopy (cryo-EM)

#### Sample preparation

Purified samples of *Sg* Enc_2-MIBS, *Sg* Enc (*E. coli BL21* (DE)- expressed), *Sg* Enc (*S. coelicolor* A3(2)-expressed), and *Sg* Enc_2-MIBS (20 mM cAMP) were concentrated to 2-3 mg/mL in 150 mM NaCl, 20 mM Tris pH 7.5 for *Sg* Enc_2-MIBS, *Sg* Enc (*E. coli BL21* (DE)-expressed), and *Sg* Enc (*S. coelicolor* A3(2)- expressed). *Sg* Enc_2-MIBS + 20 mM cAMP was prepared in buffer containing 10 mM Hepes pH 7.5, 5 mM MgCl_2_, and 20 mM cAMP. Samples were applied to freshly glow discharged Quantifoil R1.2/1.3 Cu 200 mesh grids and were immediately plunged into liquid ethane using an FEI Vitrobot Mark IV (100% humidity, 22°C, blot force 20, blot time 4 s). The grids were then clipped and stored in liquid nitrogen until data collection. *Data collection*: Cryo-EM movies of *Sg* Enc_2-MIBS and *Sg* Enc_2-MIBS (20 mM cAMP) were collected using a ThermoFisher Scientific Talos Arctica cryo-electron microscope operating at 200 kV equipped with a Gatan K2 direct electron detector. Cryo-EM movies of *Sg* Enc (*E. coli BL21* (DE)-expressed) and *Sg* Enc (*S. coelicolor* A3(2)-expressed) were collected using a ThermoFisher Scientific Glacios cryo-electron microscope operating at 200 kV equipped with a Gatan K2 direct electron detector. The Leginon^93^ software package was used to collect all datasets at a 45,000x magnification and pixel size of 0.91 Å. Further data collection statistics can be found in Supplementary Table 1.

#### Data processing

cryoSPARC v 4.4.0+231114^94^ was used to process the *Sg* Enc_2-MIBS dataset. 1,454 movies were imported, motion corrected by patch motion correction, and the contrast transfer function (CTF) fit was estimated using patch CTF estimation. Movies with a CTF fit worse than 5 Å were excluded from the dataset, resulting in 1,302 movies. Manual picker was used to select 296 particles, which were used to create templates for particle picking. 54,206 particles with a box size of 440 pixels were selected and extracted from the dataset using template picker. 2 rounds of 2D classification were then used to further sort the particles, resulting in 51,501 remaining particles. Initial volumes were created using ab-initio reconstruction with 5 classes and I symmetry, resulting in a major class containing 51,173 particles. These particles were then used for homogeneous refinement against the initial ab-initio reconstruction map with I symmetry imposed, per particle defocus optimization, per-group CTF parameterization, and Ewald sphere correction enabled with a positive curvature sign, resulting in a 3.02 Å map. The particles were then symmetry-expanded according to I symmetry and used for downstream 3D classification. For analysis of the CBDs, 3,070,380 symmetry-expanded particles were sorted by 3D classification without alignment into 10 classes using a mask containing two CBDs arranged around the two-fold axis of symmetry with a target resolution of 3.5 Å, an O-EM learning rate init of 0.75, class similarity of 0.25, and force hard classification enabled. The classes were then manually inspected, and the class with the best-resolved CBD density was reconstructed using homogeneous reconstruction only, resulting in a 3.49 Å map. The same 3D classification parameters were used for analysis of the CLD, except a map encompassing the three-fold axis of the shell and CLD density was used for classification. Particle subtraction was performed using a mask of the *Sg* Enc shell to generate shell-subtracted particles. 2D classification was then used to classify and sort shell-subtracted particles.

cryoSPARC v 4.4.0+231114^94^ was used to process the *Sg* Enc (*E. coli BL21* (DE)-expressed) dataset. 1,930 movies were imported, motion corrected with patch motion correction, and CTF fit was estimated using patch CTF estimation. Movies with CTF fit values worse than 5 Å were removed from the dataset, resulting in 1,898 remaining movies. 454 particles were manually selected from micrographs and used to generate templates for particle picking. 195,618 particles were selected using template picker and extracted with a box size of 440 pixels. Two rounds of 2D classification were then used to sort the particles, resulting in 135,962 remaining particles. An initial map was then created using ab-initio reconstruction with 3 classes and I symmetry imposed, resulting in a majority class containing 135,222 particles. These particles were then used for homogeneous refinement against the ab-initio map with I symmetry imposed, per-particle defocus optimization, per-group CTF parameterization, spherical aberration fit enabled, tetrafoil fit enabled, Ewald sphere correction enabled with a negative curvature sign, and minimize over per-particle scale enabled, resulting in a 2.75 Å resolution map. The particles were then polished using reference based motion correction, resulting in 135,027 particles which were used to perform a final homogeneous refinement with I symmetry imposed, per-particle defocus optimization, per-group CTF parameterization, spherical aberration fit enabled, tetrafoil fit enabled, Ewald sphere correction with negative curvature, and minimize over per-particle scale enabled, resulting in a 2.71 Å resolution map. The particles were then symmetry-expanded and used for subsequent 3D classification without alignment of the CBD and two-fold of the shell. For analysis of the CBDs, 8,101,620 symmetry expanded particles were sorted by 3D classification into 10 classes using a mask containing two CBDs along the two-fold axis with a target resolution of 3.5 Å, an O-EM learning rate init of 0.75, class similarity of 0.25, and force hard classification enabled. The 3D classes were then manually inspected and the class with the best resolved CBD containing 828,041 symmetry-expanded particles was then reconstructed using homogeneous reconstruction only, resulting in a 3.10 Å map. The same 3D classification parameters were used for analysis of the three-fold, using a map encompassing the three-fold shell and CLD density. Particle subtraction was performed using a mask of the *Sg* Enc shell to generate shell-subtracted particles. 2D classification was then used to classify and sort shell-subtracted particles.

cryoSPARC v 4.4.0+231114^94^ was used to process the *Sg* Enc (*S. coelicolor* A3(2)-expressed) dataset. 156 movies were imported, motion corrected by patch motion correction, and the CTF fit was estimated using patch CTF estimation. Manual picker was used to select 174 particles, which were used to create templates for particle picking. 7,003 particles were identified using template picker and extracted with a box size of 440 pixels. 2 rounds of 2D classification were used to further sort the particles, resulting in 4,119 particles. Ab-initio reconstruction was performed with I symmetry and 4 classes, resulting in a majority class containing 4,091 particles. These particles were then used to for homogeneous refinement against the ab-initio volume with I symmetry imposed, per-particle defocus optimization, per-group CTF parameterization, tetrafoil fit enabled, anisotropic magnification fit enabled, and Ewald sphere correction with negative curvature, resulting in a 3.47 Å reconstructed map. Particle subtraction was performed using a mask of the *Sg* Enc shell to generate shell-subtracted particles. 2D classification was then used to classify and sort shell-subtracted particles.

cryoSPARC v 4.4.0+231114^94^ was used to process the *Sg* Enc_2-MIBS + 20 mM cAMP dataset. 86 movies were imported, motion corrected using patch motion correction, and the CTF fit was estimated using patch CTF. Exposures with CTF fits worse than 5 Å were removed from the dataset, resulting in 83 remaining exposures. 315 particles were manually selected and used to establish templates for particle picking. 9,265 particles were identified using template picker and extracted with a box size of 440 pixels. The particles were sorted by two rounds of 2D classification, resulting in 9,181 particles. Ab-initio reconstruction was used with these particles to generate starting volumes with I symmetry imposed and two classes, resulting in a majority class containing 9,170 particles. These particles were then used for homogeneous refinement with I symmetry, per-particle defocus optimization, per-group CTF parameterization, spherical aberration fit enabled, tetrafoil fit enabled, anisotropic magnification fit enabled, Ewald sphere correction with negative curvature, resulting in a 2.92 Å map. Particles were polished using reference based motion correction, and then used for an additional homogenous refinement with I symmetry, per-particle defocus optimization, per-group CTF parameterization, spherical aberration fit, tetrafoil fit, anisotropic magnification fit, and Ewald sphere correction with a positive curvature sign, resulting in a 2.93 Å reconstructed map. The particles were then symmetry-expanded according to I symmetry and used for 3D classification. For analysis of the CBDs, 550,200 symmetry-expanded particles were sorted into by 3D classification without alignment using a mask containing two CBDs at the two-fold axis into 4 classes with a target resolution of 3.5 Å, O-EM learning rate init of 0.75, force hard classification enabled, and a class similarity of 0.25. The 3D classes were manually inspected, and the class with the best resolved CBD containing 142,420 symmetry-expanded particles was then reconstructed using homogeneous reconstruction only, resulting in a 3.43 Å reconstructed map.

#### Model building

For building the *Sg* Enc model, an AlphaFold^95^ generated model of *Sg* Enc was obtained from UniProt^96^ and manually placed in the I-refined cryo-EM map using ChimeraX v 1.5^97^, and the fit was refined using the Fit in Map command. The model was then refined iteratively by alternating rounds of manual refinement in Coot v 8.9.1^98,99^ and real-space refinement using Phenix v 1.20.1-4487^100,101^. Symmetry operators were identified from the map using the map_symmetry command and applied using the apply_ncs command to generate an icosahedral shell with 60 copies of the *Sg* Enc protomer. The NCS-expanded shell was then refined against the I-refined cryo-EM map using real-space refinement with NCS restraints. A single protomer of the NCS-refined model was then refined against the C1 map of the CBDs generated by 3D classification and homogenous reconstruction using real space refinement to improve the model of the CBD. BioMT operators were identified using the find_NCS command and manually applied to the headers of the model. Model building of *Sg* Enc_2-MIBS and *Sg* Enc_2-MIBS + 20 mM cAMP followed a nearly identical strategy, except that the *Sg* Enc model was used as a starting model.

## Data availability

The cryo-EM maps and structural models of *Sg* Enc_2-MIBS, *Sg* Enc_2-MIBS + cAMP, and *Sg* Enc have been deposited and are publicly available in the Electron Microscopy Data Bank (EMDB-44554, EMDB-44557, and EMDB-44553) and Protein Data Bank (PDB ID: 9BHV, PDB ID: 9BI0, and PDB ID: 9BHU). Sequence similarity networks and sequence alignments are supplied as supplementary data.

## Acknowledgements

We acknowledge funding from the NIH (R35GM133325). Research reported in this work was supported by the University of Michigan Cryo-EM Facility (U-M Cryo-EM). U-M Cryo-EM is grateful for support from the U-M Life Sciences Institute and the U-M Biosciences Initiative. Molecular graphics and analyses were performed using UCSF ChimeraX developed by the Resource for Biocomputing, Visualization, and Informatics at the University of California, San Francisco, with support from the National Institutes of Health R01GM129325 and the Office of Cyber Infrastructure and Computational Biology, National Institute of Allergy and Infectious Diseases. We thank Robert Benisch for the design and production of the *Ec* CAP plasmid.

## Author contributions

M.P.A. and T.W.G. designed the project. M.P.A. and T.W.G conducted the bioinformatic and phylogenetic analyses. M.P.A. conducted all laboratory experiments. M.P.A. collected cryo-EM data, and M.P.A. and T.W.G. processed and analyzed the cryo-EM data. M.P.A. built the structural models. M.P.A. and T.W.G. wrote the manuscript. T.W.G. oversaw the project in its entirety.

## References

1 Christianson, D. W. Structural and Chemical Biology of Terpenoid Cyclases. Chemical Reviews 117, 11570–11648 (2017). 10.1021/acs.chemrev.7b00287

2 Rudolf, J. D., Alsup, T. A., Xu, B. & Li, Z. Bacterial terpenome. Nat Prod Rep 38, 905–980 (2021). 10.1039/d0np00066c

3 Tholl, D. in Biotechnology of Isoprenoids (eds Jens Schrader & Jörg Bohlmann) 63-106 (Springer International Publishing, 2015).

4 Harvey, A. L. Natural products in drug discovery. Drug Discovery Today 13, 894–901 (2008). 10.1016/j.drudis.2008.07.004

5 Cane, D. E. & Ikeda, H. Exploration and mining of the bacterial terpenome. Acc Chem Res 45, 463–472 (2012). 10.1021/ar200198d

6 Dickschat, J. S. Bacterial terpene cyclases. Nat Prod Rep 33, 87–110 (2016). 10.1039/c5np00102a

7 Juttner, F. & Watson, S. B. Biochemical and ecological control of geosmin and 2-methylisoborneol in source waters. Appl Environ Microbiol 73, 4395–4406 (2007). 10.1128/AEM.02250-06

8 Cane, D. E. & Ikeda, H. Exploration and Mining of the Bacterial Terpenome. Accounts of Chemical Research 45, 463–472 (2012). 10.1021/ar200198d

9 Becher, P. G. et al. Developmentally regulated volatiles geosmin and 2-methylisoborneol attract a soil arthropod to Streptomyces bacteria promoting spore dispersal. Nat Microbiol 5, 821–829 (2020). 10.1038/s41564-020-0697-x

10 Zaroubi, L. et al. The Ubiquitous Soil Terpene Geosmin Acts as a Warning Chemical. Appl Environ Microbiol 88, e0009322 (2022). 10.1128/aem.00093-22

11 Garbeva, P., Avalos, M., Ulanova, D., van Wezel, G. P. & Dickschat, J. S. Volatile sensation: The chemical ecology of the earthy odorant geosmin. Environ Microbiol 25, 1565–1574 (2023). 10.1111/1462-2920.16381

12 Saha, M. & Fink, P. Algal volatiles - the overlooked chemical language of aquatic primary producers. Biol Rev Camb Philos Soc 97, 2162–2173 (2022). 10.1111/brv.12887

13 Wang, C. M. & Cane, D. E. Biochemistry and molecular genetics of the biosynthesis of the earthy odorant methylisoborneol in Streptomyces coelicolor. J Am Chem Soc 130, 8908–8909 (2008). 10.1021/ja803639g

14 Giglio, S., Chou, W. K., Ikeda, H., Cane, D. E. & Monis, P. T. Biosynthesis of 2-methylisoborneol in cyanobacteria. Environ Sci Technol 45, 992–998 (2011). 10.1021/es102992p

15 Wang, Z., Xu, Y., Shao, J., Wang, J. & Li, R. Genes associated with 2-methylisoborneol biosynthesis in cyanobacteria: isolation, characterization, and expression in response to light. PLoS One 6, e18665 (2011). 10.1371/journal.pone.0018665

16 Dickschat, J. S. et al. Biosynthesis of the off-flavor 2-methylisoborneol by the myxobacterium Nannocystis exedens. Angew Chem Int Ed Engl 46, 8287–8290 (2007). 10.1002/anie.200702496

17 Koksal, M., Chou, W. K., Cane, D. E. & Christianson, D. W. Structure of 2-methylisoborneol synthase from Streptomyces coelicolor and implications for the cyclization of a noncanonical C-methylated monoterpenoid substrate. Biochemistry 51, 3011–3020 (2012). 10.1021/bi201827a

18 Komatsu, M., Tsuda, M., Omura, S., Oikawa, H. & Ikeda, H. Identification and functional analysis of genes controlling biosynthesis of 2-methylisoborneol. Proc Natl Acad Sci U S A 105, 7422–7427 (2008). 10.1073/pnas.0802312105

19 Koksal, M., Chou, W. K., Cane, D. E. & Christianson, D. W. Structure of geranyl diphosphate C-methyltransferase from Streptomyces coelicolor and implications for the mechanism of isoprenoid modification. Biochemistry 51, 3003–3010 (2012). 10.1021/bi300109c

20 Kawamoto, S. et al. Molecular and functional analyses of the gene (eshA) encoding the 52-kilodalton protein of Streptomyces coelicolor A3(2) required for antibiotic production. J Bacteriol 183, 6009–6016 (2001). 10.1128/JB.183.20.6009-6016.2001

21 Passner, J. M., Schultz, S. C. & Steitz, T. A. Modeling the cAMP-induced Allosteric Transition Using the Crystal Structure of CAP-cAMP at 2.1Å Resolution. Journal of Molecular Biology 304, 847–859 (2000). 10.1006/jmbi.2000.4231

22 Andreas, M. P. & Giessen, T. W. Large-scale computational discovery and analysis of virus-derived microbial nanocompartments. Nat Commun 12, 4748 (2021). 10.1038/s41467-021-25071-y

23 Nichols, R. J. et al. Discovery and characterization of a novel family of prokaryotic nanocompartments involved in sulfur metabolism. Elife 10 (2021). 10.7554/eLife.59288

24 Giessen, T. W. Encapsulins. Annu Rev Biochem (2022). 10.1146/annurev-biochem-040320-102858

25 Nichols, R. J., Cassidy-Amstutz, C., Chaijarasphong, T. & Savage, D. F. Encapsulins: molecular biology of the shell. Crit Rev Biochem Mol Biol 52, 583–594 (2017). 10.1080/10409238.2017.1337709

26 Sutter, M. et al. Structural basis of enzyme encapsulation into a bacterial nanocompartment. Nat Struct Mol Biol 15, 939–947 (2008). 10.1038/nsmb.1473

27 Giessen, T. W. & Silver, P. A. Widespread distribution of encapsulin nanocompartments reveals functional diversity. Nat Microbiol 2, 17029 (2017). 10.1038/nmicrobiol.2017.29

28 Heinemann, J. et al. Fossil record of an archaeal HK97-like provirus. Virology 417, 362–368 (2011). 10.1016/j.virol.2011.06.019

29 Altenburg, W. J., Rollins, N., Silver, P. A. & Giessen, T. W. Exploring targeting peptide-shell interactions in encapsulin nanocompartments. Sci Rep 11, 4951 (2021). 10.1038/s41598-021-84329-z

30 Sutter, M. et al. Structural basis of enzyme encapsulation into a bacterial nanocompartment. Nat Struct Mol Biol 15, 939–947 (2008). 10.1038/nsmb.1473

31 McHugh, C. A. et al. A virus capsid-like nanocompartment that stores iron and protects bacteria from oxidative stress. EMBO J 33, 1896–1911 (2014). 10.15252/embj.201488566

32 Giessen, T. W. et al. Large protein organelles form a new iron sequestration system with high storage capacity. Elife 8 (2019). 10.7554/eLife.46070

33 Akita, F. et al. The crystal structure of a virus-like particle from the hyperthermophilic archaeon Pyrococcus furiosus provides insight into the evolution of viruses. J Mol Biol 368, 1469–1483 (2007). 10.1016/j.jmb.2007.02.075

34 Kwon, S., Andreas, M. P. & Giessen, T. W. Structure and heterogeneity of a highly cargo-loaded encapsulin shell. J Struct Biol 215, 108022 (2023). 10.1016/j.jsb.2023.108022

35 Eren, E. et al. Structural characterization of the Myxococcus xanthus encapsulin and ferritin-like cargo system gives insight into its iron storage mechanism. Structure 30, 551–563 e554 (2022). 10.1016/j.str.2022.01.008

36 Jones, J. A., Andreas, M. P. & Giessen, T. W. Structural basis for peroxidase encapsulation inside the encapsulin from the Gram-negative pathogen Klebsiella pneumoniae. Nat Commun 15, 2558 (2024). 10.1038/s41467-024-46880-x

37 Sugano, Y. & Yoshida, T. DyP-Type Peroxidases: Recent Advances and Perspectives. Int J Mol Sci 22 (2021). 10.3390/ijms22115556

38 Almeida, A. V. et al. Condensation and Protection of DNA by the Myxococcus xanthus Encapsulin: A Novel Function. Int J Mol Sci 23 (2022). 10.3390/ijms23147829

39 Benisch, R., Andreas, M. P. & Giessen, T. W. A widespread bacterial protein compartment sequesters and stores elemental sulfur. Science Advances 10, eadk9345 (2024). doi:10.1126/sciadv.adk9345

40 Kwak, J., McCue, L. A., Trczianka, K. & Kendrick, K. E. Identification and characterization of a developmentally regulated protein, EshA, required for sporogenic hyphal branches in Streptomyces griseus. J Bacteriol 183, 3004–3015 (2001). 10.1128/JB.183.10.3004-3015.2001

41 Saito, N., Matsubara, K., Watanabe, M., Kato, F. & Ochi, K. Genetic and biochemical characterization of EshA, a protein that forms large multimers and affects developmental processes in Streptomyces griseus. J Biol Chem 278, 5902–5911 (2003). 10.1074/jbc.M208564200

42 Saito, N. et al. EshA accentuates ppGpp accumulation and is conditionally required for antibiotic production in Streptomyces coelicolor A3(2). J Bacteriol 188, 4952–4961 (2006). 10.1128/JB.00343-06

43 Jones, J. A., Benisch, R. & Giessen, T. W. Encapsulin cargo loading: progress and potential. J Mater Chem B 11, 4377–4388 (2023). 10.1039/d3tb00288h

44 Berman, H. M. et al. The cAMP binding domain: an ancient signaling module. Proc Natl Acad Sci U S A 102, 45–50 (2005). 10.1073/pnas.0408579102

45 Kannan, N., et al. Evolution of allostery in the cyclic nucleotide binding module. Genome Biol 8, R264 (2007). 10.1186/gb-2007-8-12-r264

46 Krol, E., Werel, L., Essen, L. O. & Becker, A. Structural and functional diversity of bacterial cyclic nucleotide perception by CRP proteins. microLife 4 (2023). 10.1093/femsml/uqad024

47 Popovych, N., Tzeng, S.-R., Tonelli, M., Ebright, R. H. & Kalodimos, C. G. Structural basis for cAMP-mediated allosteric control of the catabolite activator protein. Proceedings of the National Academy of Sciences 106, 6927–6932 (2009). doi:10.1073/pnas.0900595106

48 Yariv, B. et al. Using evolutionary data to make sense of macromolecules with a "face-lifted" ConSurf. Protein Sci 32, e4582 (2023). 10.1002/pro.4582

49 Ashkenazy, H., et al. ConSurf 2016: an improved methodology to estimate and visualize evolutionary conservation in macromolecules. Nucleic Acids Res 44, W344–350 (2016). 10.1093/nar/gkw408

50 Ashkenazy, H., Erez, E., Martz, E., Pupko, T. & Ben-Tal, N. ConSurf 2010: calculating evolutionary conservation in sequence and structure of proteins and nucleic acids. Nucleic Acids Res 38, W529–533 (2010). 10.1093/nar/gkq399

51 Belduz, A. O., Lee, E. J. & Harman, J. G. Mutagenesis of the cyclic AMP receptor protein of Escherichia coli: targeting positions 72 and 82 of the cyclic nucleotide binding pocket. Nucleic Acids Res 21, 1827–1835 (1993). 10.1093/nar/21.8.1827

52 Gronenborn, A. M., Sandulache, R., Gärtner, S. & Clore, G. M. Mutations in the cyclic AMP binding site of the cyclic AMP receptor protein of Escherichia coli. Biochem J 253, 801–807 (1988). 10.1042/bj2530801

53 Moore, J., Kantorow, M., Vanderzwaag, D. & McKenney, K. Escherichia coli cyclic AMP receptor protein mutants provide evidence for ligand contacts important in activation. J Bacteriol 174, 8030–8035 (1992). 10.1128/jb.174.24.8030-8035.1992

54 Lanzilotta, W. N. et al. Structure of the CO sensing transcription activator CooA. Nature Structural Biology 7, 876–880 (2000). 10.1038/82820

55 Joyce, M. G. et al. CprK Crystal Structures Reveal Mechanism for Transcriptional Control of Halorespiration *. Journal of Biological Chemistry 281, 28318–28325 (2006). 10.1074/jbc.M602654200

56 Zhao, M.-X. et al. Structural basis for the allosteric control of the global transcription factor NtcA by the nitrogen starvation signal 2-oxoglutarate. Proceedings of the National Academy of Sciences 107, 12487–12492 (2010). doi:10.1073/pnas.1001556107

57 Brelidze, T. I., Carlson, A. E., Sankaran, B. & Zagotta, W. N. Structure of the carboxy-terminal region of a KCNH channel. Nature 481, 530–533 (2012). 10.1038/nature10735

58 Marques-Carvalho, M. J., et al. Structural, Biochemical, and Functional Characterization of the Cyclic Nucleotide Binding Homology Domain from the Mouse EAG1 Potassium Channel. Journal of Molecular Biology 423, 34–46 (2012). 10.1016/j.jmb.2012.06.025

59 Haitin, Y., Carlson, A. E. & Zagotta, W. N. The structural mechanism of KCNH-channel regulation by the eag domain. Nature 501, 444–448 (2013). 10.1038/nature12487

60 van Kempen, M. et al. Fast and accurate protein structure search with Foldseek. Nature Biotechnology (2023). 10.1038/s41587-023-01773-0

61 Holm, L., Laiho, A., Törönen, P. & Salgado, M. DALI shines a light on remote homologs: One hundred discoveries. Protein Science 32, e4519 (2023). 10.1002/pro.4519

62 Gabler, F. et al. Protein Sequence Analysis Using the MPI Bioinformatics Toolkit. Curr Protoc Bioinformatics 72, e108 (2020). 10.1002/cpbi.108

63 Zimmermann, L. et al. A Completely Reimplemented MPI Bioinformatics Toolkit with a New HHpred Server at its Core. J Mol Biol 430, 2237–2243 (2018). 10.1016/j.jmb.2017.12.007

64 Mirdita, M. et al. ColabFold: making protein folding accessible to all. Nature Methods 19, 679–682 (2022). 10.1038/s41592-022-01488-1

65 Cassidy-Amstutz, C. et al. Identification of a Minimal Peptide Tag for in Vivo and in Vitro Loading of Encapsulin. Biochemistry 55, 3461–3468 (2016). 10.1021/acs.biochem.6b00294

66 Faylo, J. L. et al. Structural insight on assembly-line catalysis in terpene biosynthesis. Nat Commun 12, 3487 (2021). 10.1038/s41467-021-23589-9

67 Ronnebaum, T. A., Eaton, S. A., Brackhahn, E. A. E. & Christianson, D. W. Engineering the Prenyltransferase Domain of a Bifunctional Assembly-Line Terpene Synthase. Biochemistry 60, 3162–3172 (2021). 10.1021/acs.biochem.1c00600

68 Ronnebaum, T. A., Gupta, K. & Christianson, D. W. Higher-order oligomerization of a chimeric αβγ bifunctional diterpene synthase with prenyltransferase and class II cyclase activities is concentration-dependent. Journal of Structural Biology 210, 107463 (2020). 10.1016/j.jsb.2020.107463

69 Gao, K., Oerlemans, R. & Groves, M. R. Theory and applications of differential scanning fluorimetry in early-stage drug discovery. Biophys Rev 12, 85–104 (2020). 10.1007/s12551-020-00619-2

70 Dremier, S., Kopperud, R., Doskeland, S. O., Dumont, J. E. & Maenhaut, C. Search for new cyclic AMP-binding proteins. FEBS Letters 546, 103–107 (2003). 10.1016/S0014-5793(03)00561-1

71 Bertinetti, D. et al. Chemical tools selectively target components of the PKA system. BMC Chemical Biology 9, 3 (2009). 10.1186/1472-6769-9-3

72 Endoh, T. & Engel Joanne, N. CbpA: a Polarly Localized Novel Cyclic AMP-Binding Protein in Pseudomonas aeruginosa. Journal of Bacteriology 191, 7193–7205 (2009). 10.1128/jb.00970-09

73 Jäger, A. V. et al. Identification of novel cyclic nucleotide binding proteins in Trypanosoma cruzi. Molecular and Biochemical Parasitology 198, 104–112 (2014). 10.1016/j.molbiopara.2015.02.002

74 Jones, J. A., Andreas, M. P. & Giessen, T. W. Exploring the Extreme Acid Tolerance of a Dynamic Protein Nanocage. Biomacromolecules 24, 1388–1399 (2023). 10.1021/acs.biomac.2c01424

75 Ross, J. et al. Pore dynamics and asymmetric cargo loading in an encapsulin nanocompartment. Sci Adv 8, eabj4461 (2022). 10.1126/sciadv.abj4461

76 Adamson, L. S. R. et al. Pore structure controls stability and molecular flux in engineered protein cages. Sci Adv 8, eabl7346 (2022). 10.1126/sciadv.abl7346

77 Joyce, M. G. CprK Crystal Structures Reveal Mechanism for Transcriptional Control of Halorespiration*. (2006).

78 Ober, V. T. et al. Purine nucleosides replace cAMP in allosteric regulation of PKA in trypanosomatid pathogens. eLife 12, RP91040 (2024). 10.7554/eLife.91040

79 Bachmaier, S. et al. Nucleoside analogue activators of cyclic AMP-independent protein kinase A of Trypanosoma. Nature Communications 10, 1421 (2019). 10.1038/s41467-019-09338-z

80 Zallot, R., Oberg, N. & Gerlt, J. A. The EFI Web Resource for Genomic Enzymology Tools: Leveraging Protein, Genome, and Metagenome Databases to Discover Novel Enzymes and Metabolic Pathways. Biochemistry 58, 4169–4182 (2019). 10.1021/acs.biochem.9b00735

81 Oberg, N., Zallot, R. & Gerlt, J. A. EFI-EST, EFI-GNT, and EFI-CGFP: Enzyme Function Initiative (EFI) Web Resource for Genomic Enzymology Tools. Journal of Molecular Biology (2023). 10.1016/j.jmb.2023.168018

82 Lemoine, F., et al. NGPhylogeny.fr: new generation phylogenetic services for non-specialists. Nucleic Acids Res 47, W260-W265 (2019). 10.1093/nar/gkz303

83 Katoh, K. & Standley, D. M. MAFFT Multiple Sequence Alignment Software Version 7: Improvements in Performance and Usability. Molecular Biology and Evolution 30, 772–780 (2013). 10.1093/molbev/mst010

84 Price, M. N., Dehal, P. S. & Arkin, A. P. FastTree 2 – Approximately Maximum-Likelihood Trees for Large Alignments. PLOS ONE 5, e9490 (2010). 10.1371/journal.pone.0009490

85 Price, M. N., Dehal, P. S. & Arkin, A. P. FastTree: Computing Large Minimum Evolution Trees with Profiles instead of a Distance Matrix. Molecular Biology and Evolution 26, 1641–1650 (2009). 10.1093/molbev/msp077

86 Lemoine, F. et al. Renewing Felsenstein’s phylogenetic bootstrap in the era of big data. Nature 556, 452–456 (2018). 10.1038/s41586-018-0043-0

87 Letunic, I. & Bork, P. Interactive Tree Of Life (iTOL) v5: an online tool for phylogenetic tree display and annotation. Nucleic Acids Res 49, W293–W296 (2021). 10.1093/nar/gkab301

88 Shannon, P. et al. Cytoscape: a software environment for integrated models of biomolecular interaction networks. Genome Res 13, 2498–2504 (2003). 10.1101/gr.1239303

89 Madeira, F. et al. Search and sequence analysis tools services from EMBL-EBI in 2022. Nucleic acids research 50, W276–W279 (2022). 10.1093/nar/gkac240

90 Andreas, M. P. & Giessen, T. W. Heterologous expression and purification of encapsulins in Streptomyces coelicolor. MethodsX 9, 101787 (2022). 10.1016/j.mex.2022.101787

91 Andreas, M. P. & Giessen, T. W. Cyclodipeptide oxidase is an enzyme filament. bioRxiv, 2023.2009.2025.559410 (2023). 10.1101/2023.09.25.559410

92 Schneider, C. A., Rasband, W. S. & Eliceiri, K. W. NIH Image to ImageJ: 25 years of image analysis. Nat Methods 9, 671–675 (2012).

93 Suloway, C. et al. Automated molecular microscopy: the new Leginon system. J Struct Biol 151, 41–60 (2005). 10.1016/j.jsb.2005.03.010

94 Punjani, A., Rubinstein, J. L., Fleet, D. J. & Brubaker, M. A. cryoSPARC: algorithms for rapid unsupervised cryo-EM structure determination. Nat Methods 14, 290–296 (2017). 10.1038/nmeth.4169

95 Jumper, J. et al. Highly accurate protein structure prediction with AlphaFold. Nature 596, 583–589 (2021). 10.1038/s41586-021-03819-2

96 UniProt, C. UniProt: the Universal Protein Knowledgebase in 2023. Nucleic Acids Res 51, D523–D531 (2023). 10.1093/nar/gkac1052

97 Pettersen, E. F. et al. UCSF ChimeraX: Structure visualization for researchers, educators, and developers. Protein Science 30, 70–82 (2021). 10.1002/pro.3943

98 Emsley, P., Lohkamp, B., Scott, W. G. & Cowtan, K. Features and development of Coot. Acta Crystallographica Section D: Biological Crystallography 66, 486–501 (2010). 10.1107/S0907444910007493

99 Emsley, P. & Cowtan, K. Coot: model-building tools for molecular graphics. Acta Crystallogr D Biol Crystallogr 60, 2126–2132 (2004). 10.1107/S0907444904019158

100 Liebschner, D. et al. Macromolecular structure determination using X-rays, neutrons and electrons: recent developments in Phenix. Acta Crystallogr D Struct Biol 75, 861–877 (2019). 10.1107/S2059798319011471

101 Adams, P. D. et al. PHENIX: a comprehensive Python-based system for macromolecular structure solution. Acta Crystallogr D Biol Crystallogr 66, 213–221 (2010). 10.1107/S0907444909052925

