## Supplementary Information for "The biosynthesis of the odorant 2-methylisoborneol is compartmentalized inside a protein shell"

**Table of Contents**


| | |
|---|---|
| Supplementary Fig. 1. Sequence similarity network analysis of 2-MIBSs | 3 |
| Supplementary Fig. 2 Cryo-EM workflow for *Sc*-expressed *Sg* Enc_2-MIBS | 4 |
| Supplementary Fig. 3. Cryo-EM workflow for *Ec*-expressed *Sg* Enc | 5 |
| Supplementary Fig. 4. Structural comparison between *Sg* Enc and Family 2A encapsulins | 6 |
| Supplementary Fig. 5. Unmodeled segments of *Sg* Enc | 7 |
| Supplementary Fig. 6. Cryo-EM workflow for *Sc*-expressed *Sg* Enc | 8 |
| Supplementary Fig. 7. 3D classification of cargo-loaded *Sg* Enc_2-MIBS CBDs | 9 |
| Supplementary Fig. 8. 3D classification of *Sg* Enc CBDs | 10 |
| Supplementary Fig. 9. *Sg* Enc sequence conservation | 11 |
| Supplementary Fig. 10. 3D classification of the three-fold axis of symmetry | 12 |
| Supplementary Fig. 11. Native PAGE gels for cargo-loading analysis | 13 |
| Supplementary Fig. 12. Saturation kinetics curves for 2-MIBS activity assays | 14 |
| Supplementary Fig. 13. Additional GC-MS data for *Sg* Enc_2-MIBS, *Sg* Enc, and *Sg* 2-MIBS | 15 |
| Supplementary Fig. 14. Cryo-EM workflow for *Sg* Enc_2-MIBS + 20 mM cAMP | 16 |
| Supplementary Table 1. Cryo-EM data collection, refinement, and validation statistics | 17 |
| Supplementary Table 2. Amino acid sequences of proteins used in this study | 18 |
| Supplementary References | 24 |

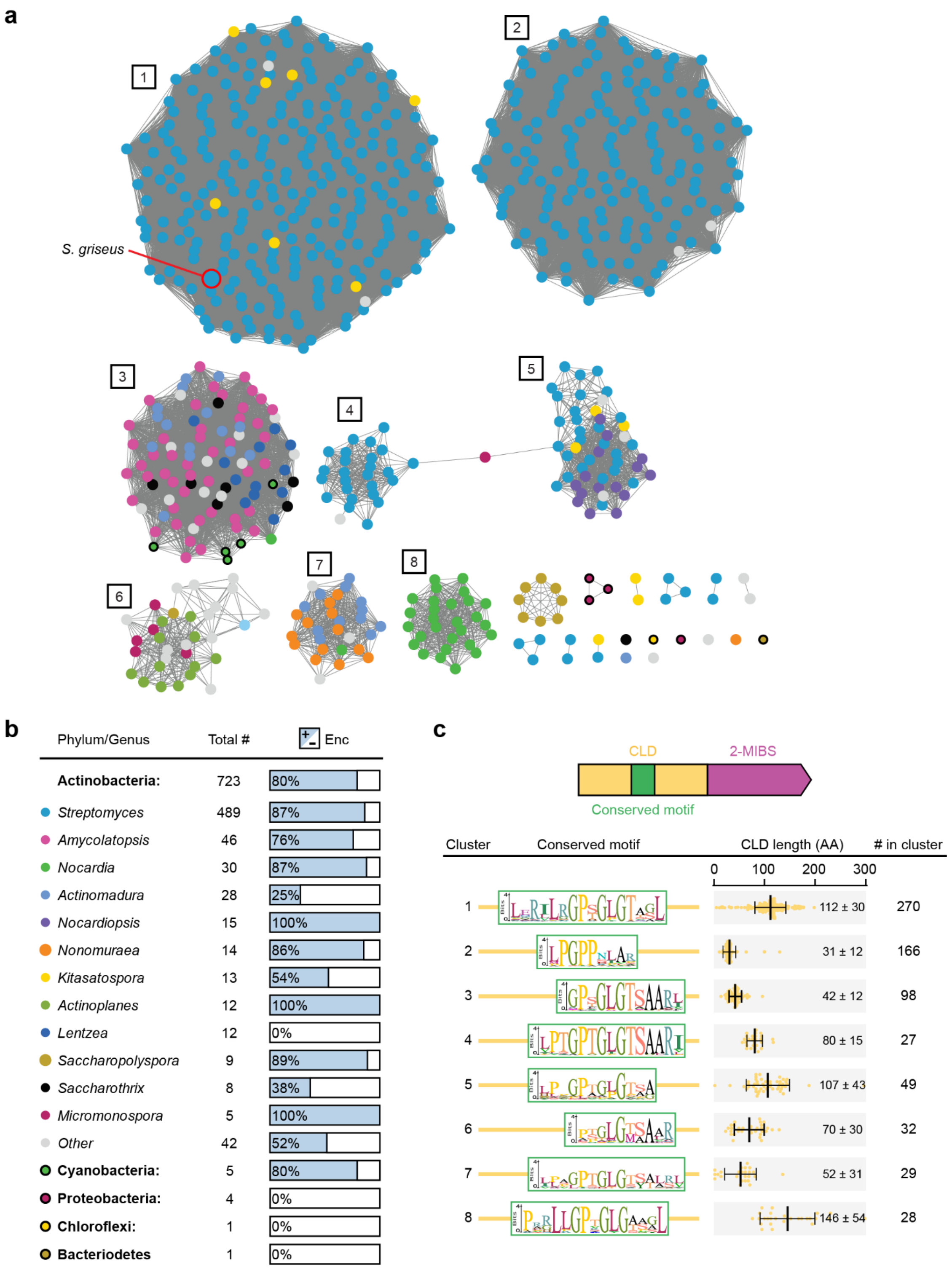


**Supplementary Fig. 1. Sequence similarity network (SSN) analysis of 2-MIBSs. a**, Sequence similarity network of all 2-MIBS sequences identified in this study. **b,** Distribution of the phyla and genera represented in the sequence similarity network. **c,** Conserved CLD peptide sequence logos and CLD lengths found within each identified SSN cluster.

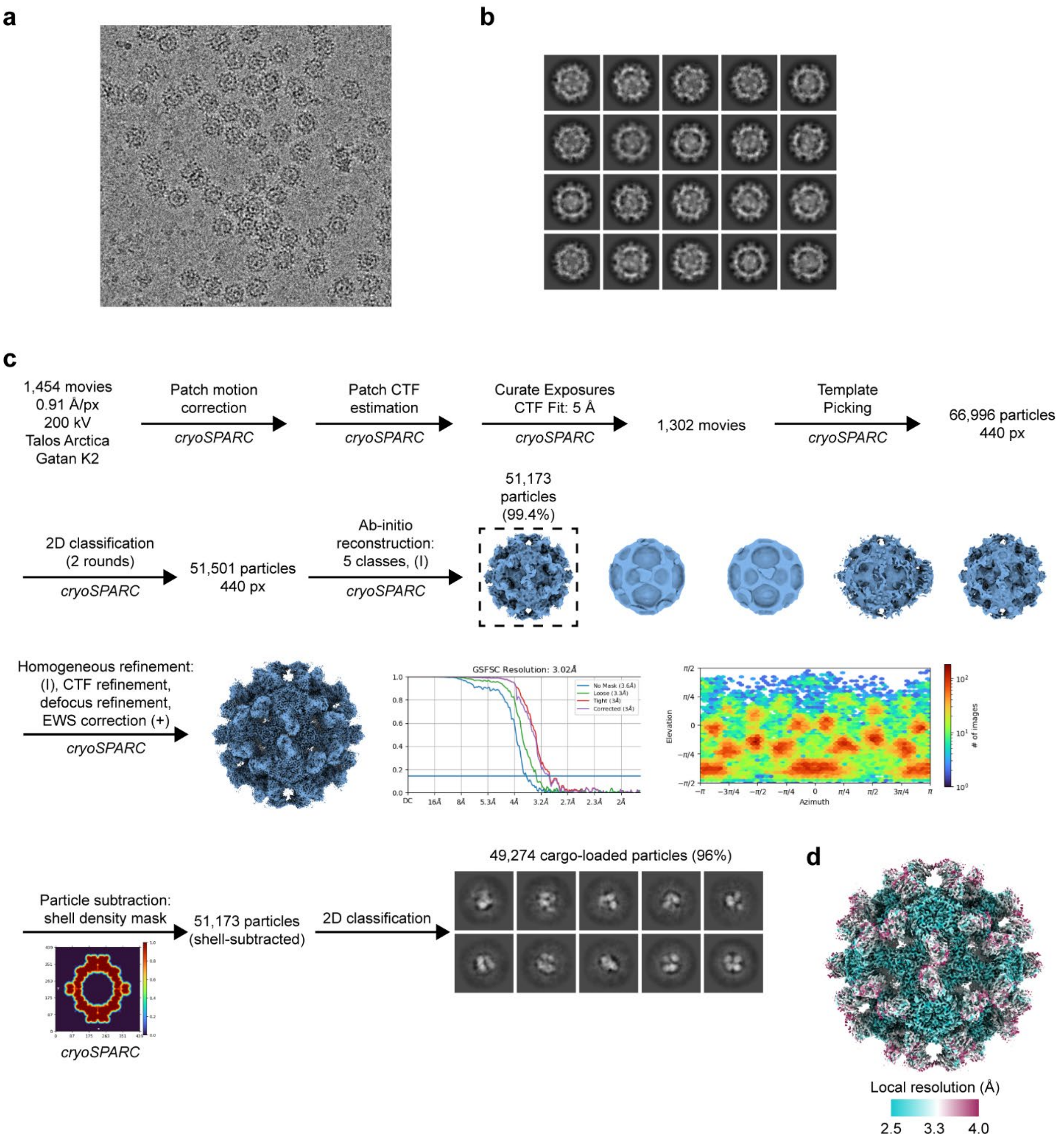


**Supplementary Fig. 2. Cryo-EM workflow for *Streptomyces coelicolor* A3(2)-expressed *Sg* Enc_2-MIBS. a,** Representative cryo-EM micrograph of *Sg* Enc_2-MIBS. **b,** Representative 2D class averages of *Sg* Enc_2-MIBS. **c,** Cryo-EM workflow for *Sg* Enc_2-MIBS. **d,** Local resolution analysis of *Sg* Enc-2-MIBS.

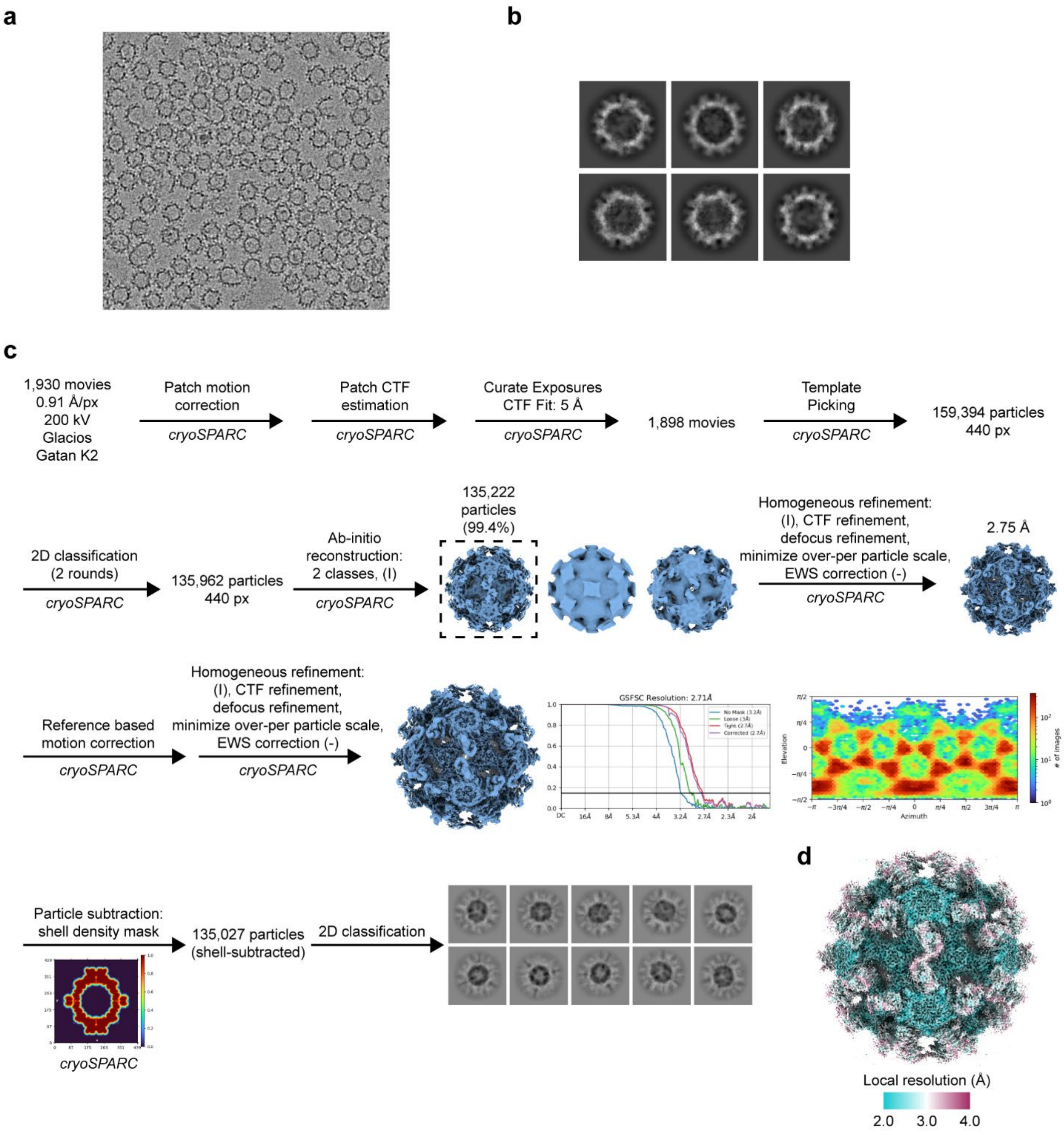


**Supplementary Fig. 3. Cryo-EM workflow for *E. coli*-expressed *Sg* Enc. a,** Representative cryo-EM micrograph of *Sg* Enc. **b,** Representative 2D class averages of *Sg* Enc. **c,** Cryo-EM workflow for *Sg* Enc. **d,** Local resolution analysis of *Sg* Enc.

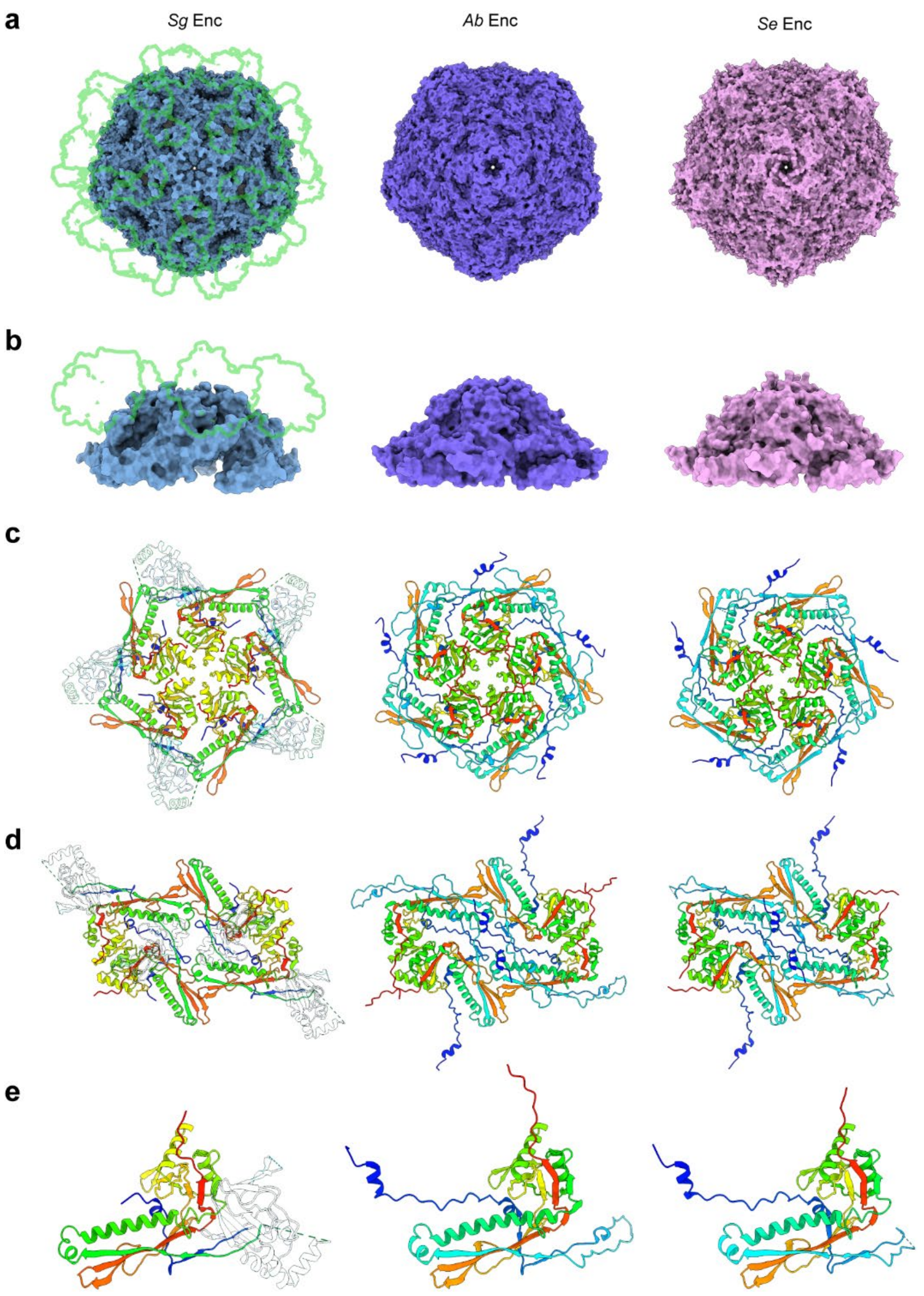


**Supplementary Fig. 4. Structural comparison between *Sg* Enc and Family 2A encapsulins.** **a**, Comparison of the Family 2B *Sg* Enc shell with the *A. baumannii* (*Ab*) (PDB ID: 8T6R)[1] and *S. elongatus* (*Se*) (PDB ID: 6X8M)[2] Family 2A encapsulin shells, centered on the five-fold axis of symmetry. Green outlines on the *Sg* Enc shell represent the CBD domains, not present in Family 2A shells. **b**, Side view of the five-fold axis of each shell, highlighting the turret-like assembly. **c**, Cartoon representation of the five-fold pentamer of each shell. Models are colored according to amino acid number (rainbow). **d**, View down the two-fold axis of symmetry showing the different orientations of the N-arms between the *Sg* Enc shell and Family 2A shells. **e**, Protomer comparison highlighting the HK97-like domain similarities between Family 2B (*Sg*) and Family 2A (*Ab*, *Se*).

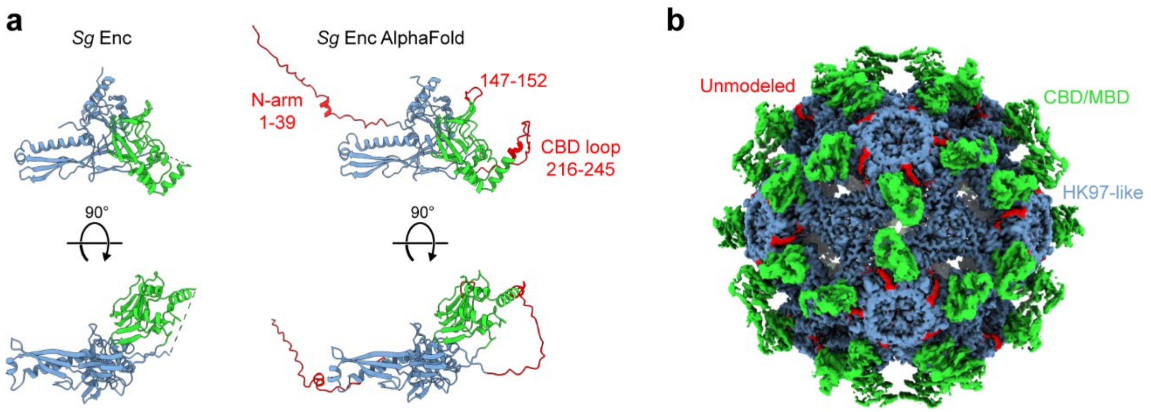


**Supplementary Fig. 5. Unmodeled segments of *Sg* Enc.** **a**, Structural comparison of the experimentally determined model of the *Sg* Enc protomer compared to an AlphaFold[3] predicted model, with red features highlighting components of *Sg* Enc that could not be modelled into the cryo-EM density map. **b**, Representative cryo-EM map of the *Sg* Enc shell highlighting observed unmodeled densities in red. Green density corresponds to CBDs (and MBDs) and blue density corresponds to the HK97-domain of *Sg* Enc.

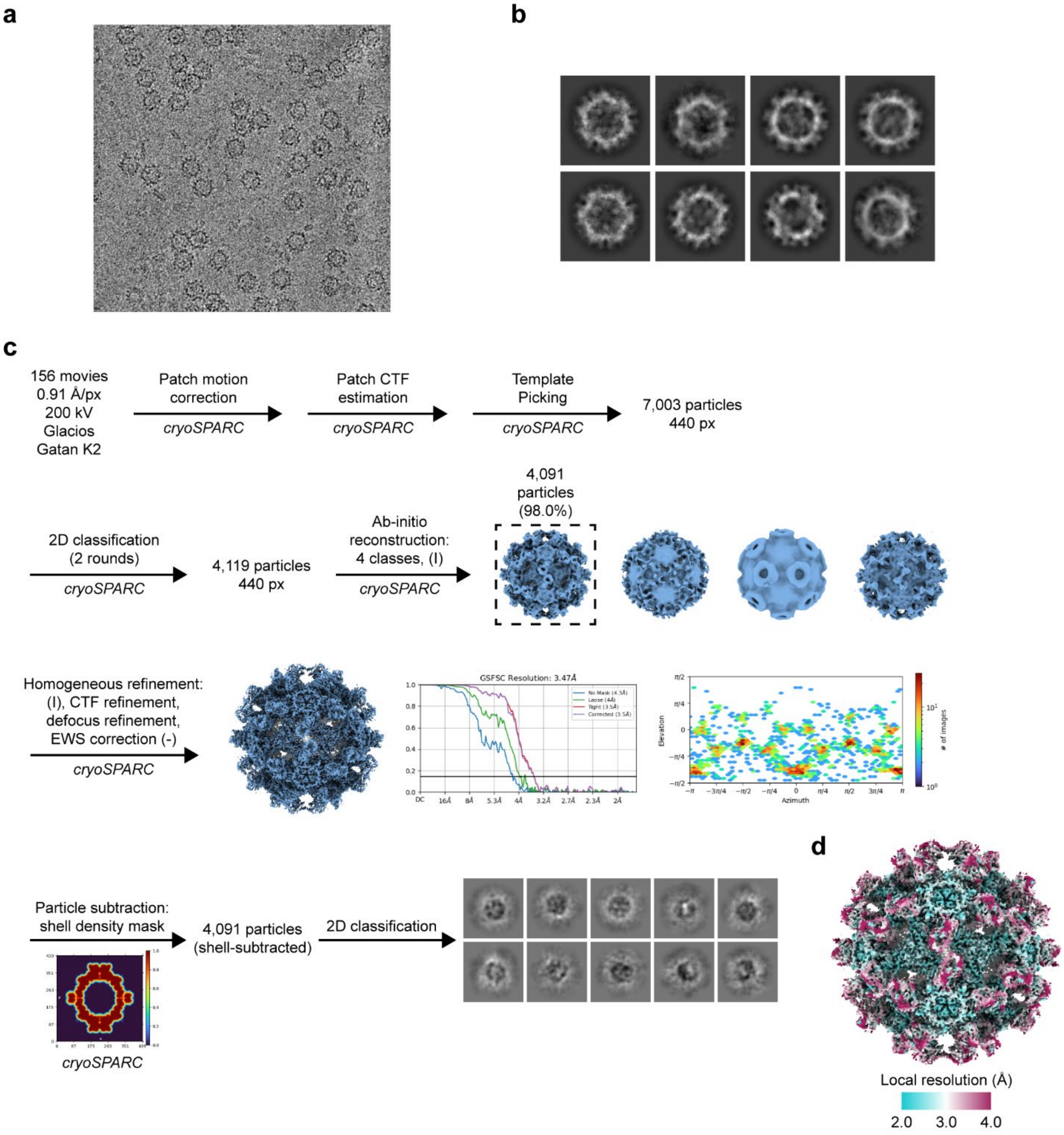


**Supplementary Fig. 6. Cryo-EM workflow for *Streptomyces coelicolor* A3(2)-expressed *Sg* Enc. a**, Representative cryo-EM micrograph of *Sg* Enc. **b**, Representative 2D class averages of *Sg* Enc. **c**, Cryo-EM workflow for *Sg* Enc. **d**, Local resolution analysis of *Sg* Enc.

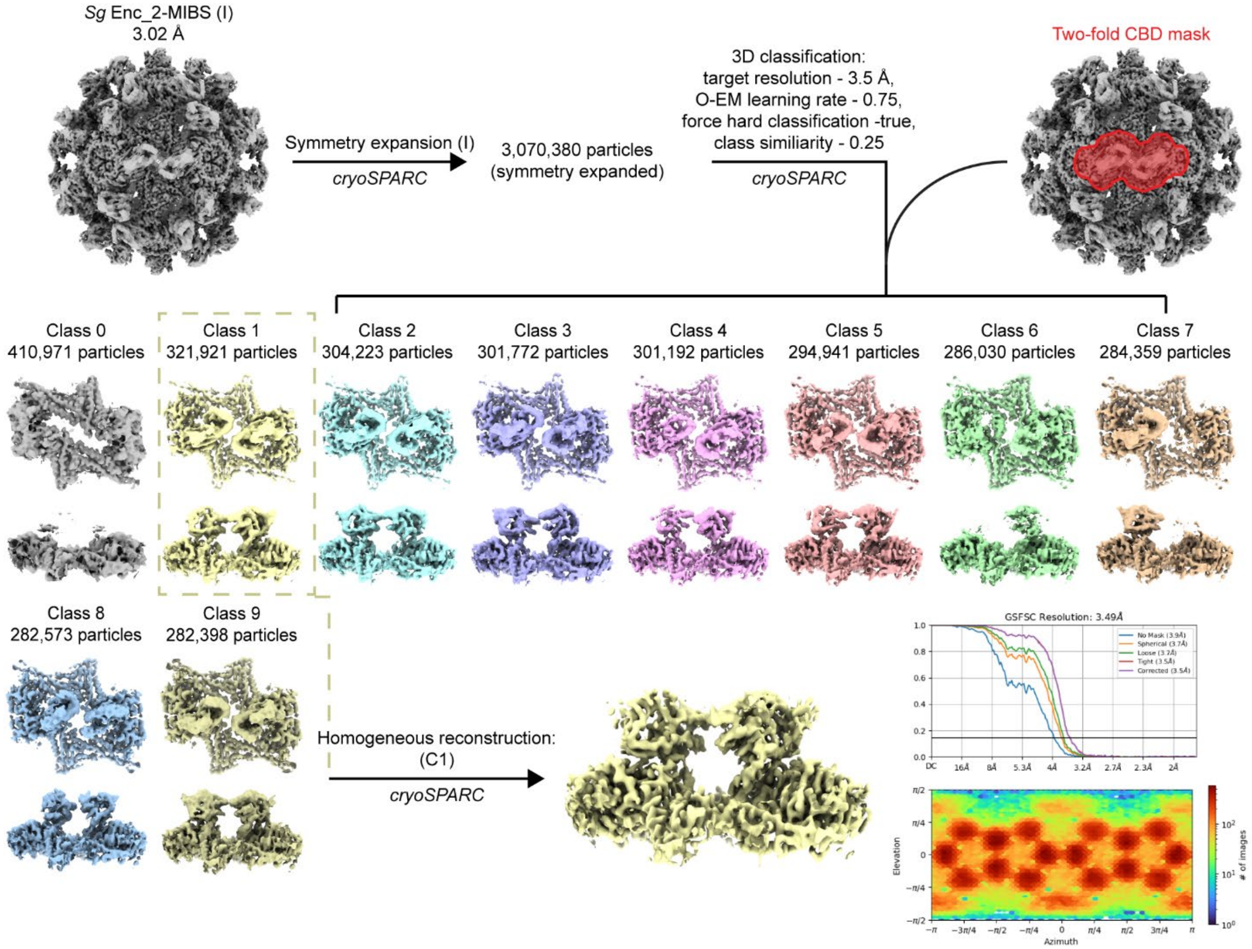


**Supplementary Fig. 7. 3D classification of cargo-loaded *Sg* Enc_2-MIBS CBDs.** Processing workflow for 3D classification followed by homogeneous reconstruction to generate an improved map of the *Sg* Enc_2-MIBS CBDs arranged around the two-fold axis of symmetry.

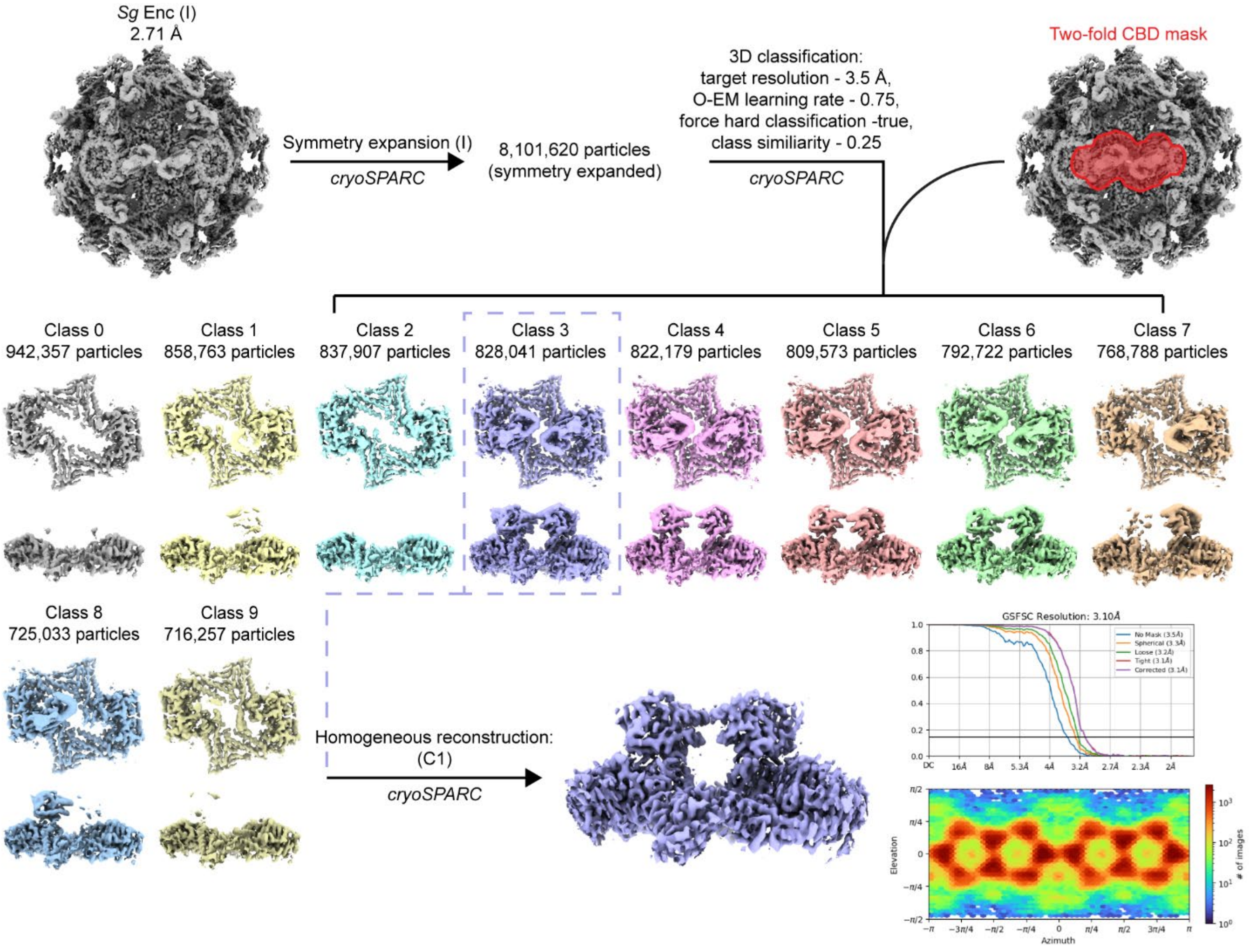


**Supplementary Fig. 8. 3D classification of *Sg* Enc CBDs.** Processing workflow for 3D classification followed by homogeneous reconstruction to generate an improved map of the *Sg* Enc CBDs arranged around the two-fold axis of symmetry.

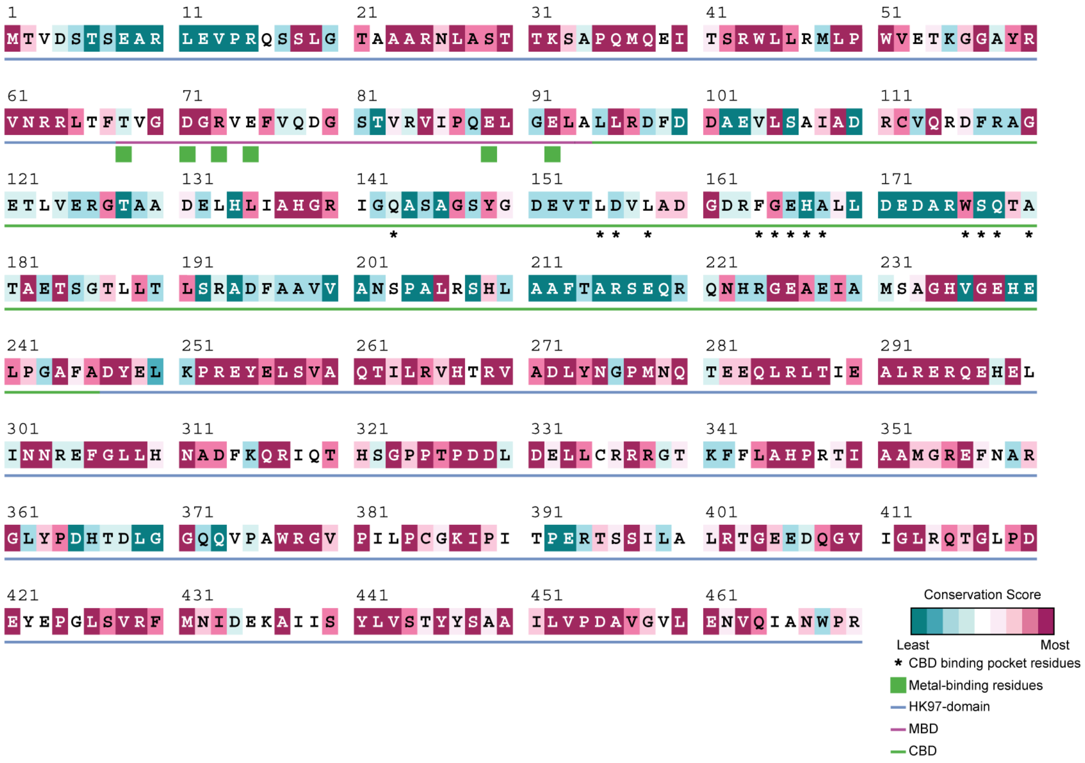

**Supplementary Fig. 9. *Sg* Enc sequence conservation.** The sequence of Sg Enc colored by the sequence conservation as computed by ConSurf[4] analysis of 600 Family 2B encapsulins is shown. Structural features of *Sg* Enc are highlighted according to the legend (bottom right).

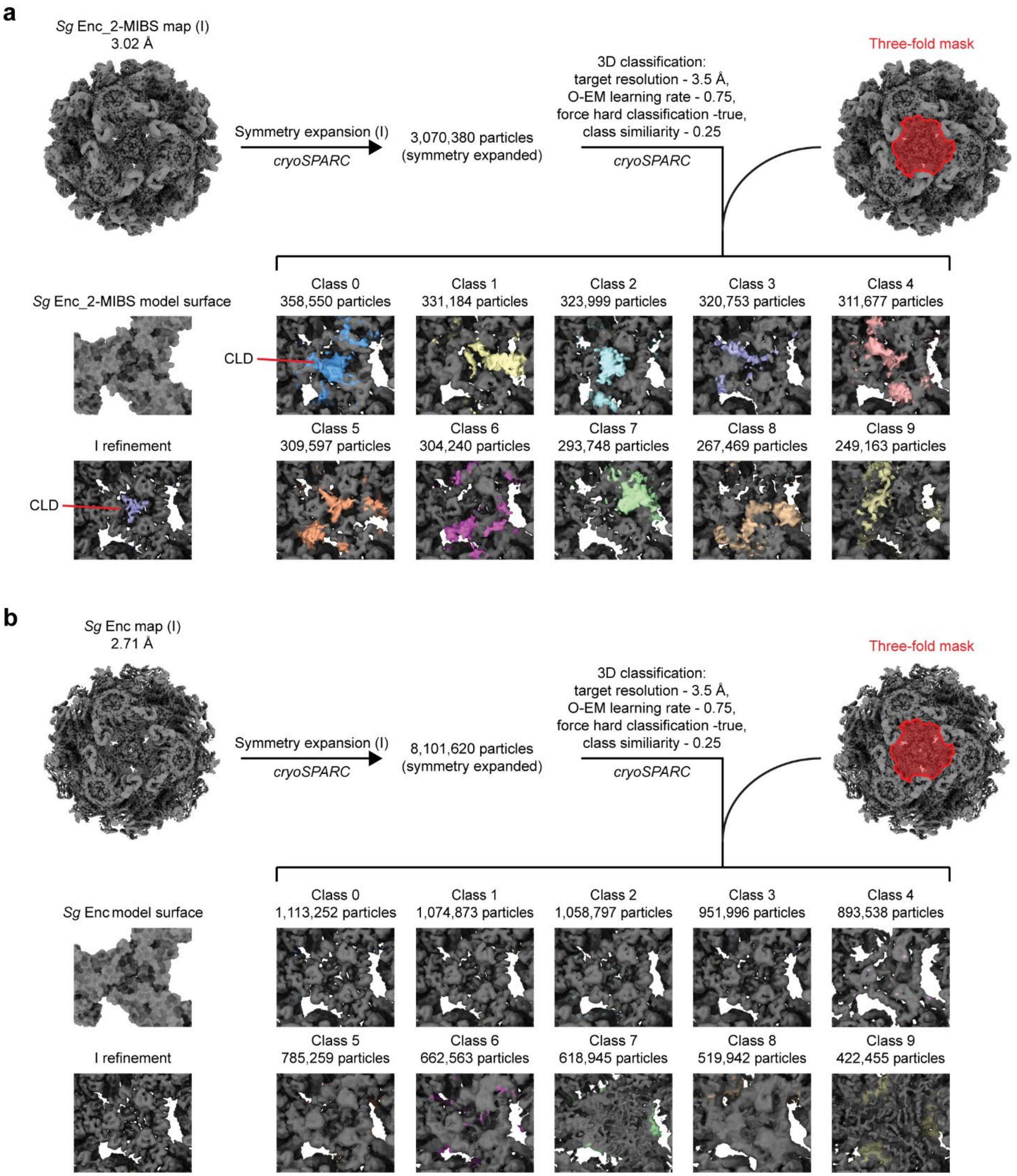


**Supplementary Fig. 10. 3D classification of the three-fold axis of symmetry of *Sg* Enc_2-MIBS and *Sg* Enc. a**, 3D classification workflow to identify CLD fragments bound to the interior of the *Sg* Enc_2-MIBS shell. **b**, 3D classification workflow highlighting that no CLD density is present at the three-fold of the *Sg* Enc shell.

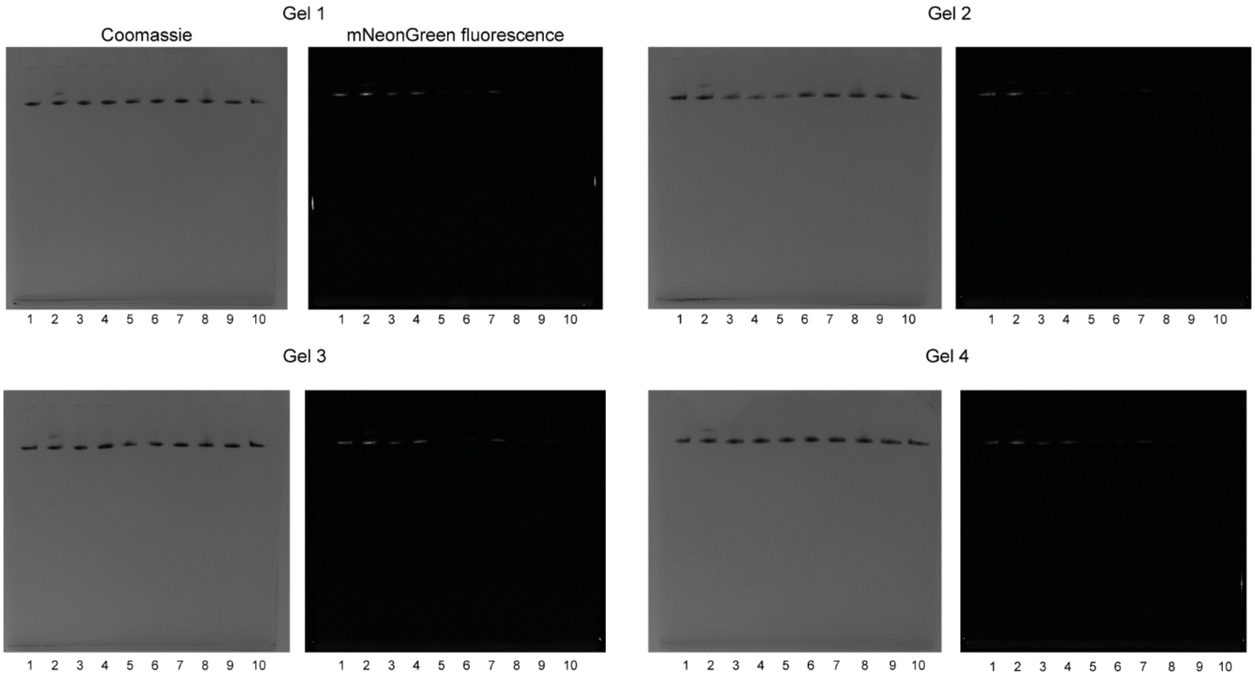


**Supplementary Fig. 11. Native PAGE gels for cargo-loading analysis.** Full native PAGE gels are shown with Coomassie stain or mNeonGreen fluorescence. Lanes are numbered according to the *Sg* Enc_CLD-mNeonGreen truncation numbering in Fig. 4f. Four replicates are shown (Gel 1 to 4).

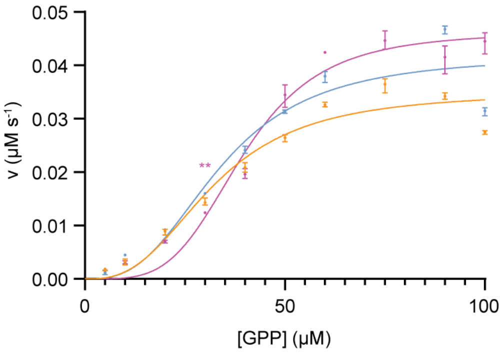


| Name | $k_{cat}$ ($s^{-1}$) | $K_m$ (μM) | $k_{cat}/K_m$ ($M^{-1}s^{-1}$) |
|---|---|---|---|
| *Sg* Enc_2-MIBS | 0.084 ± 0.006 | 33.81 ± 2.79 | 2472 ± 127.1 |
| *Sg* Enc_2-MIBS + 500 μM CAMP | 0.070 ± 0.004 | 32.15 ± 2.18 | 2177 ± 94.4 |
| *Sg* 2-MIBS | 0.092 ± 0.003 | 39.48 ± 1.37 | 2343 ± 68.2 |

**Supplementary Fig. 12. Saturation kinetics curves for 2-MIBS in vitro activity assays.** Saturation kinetics curves of *Sg* Enc_2-MIBS (blue), *Sg* Enc_2-MIBS + 500 μM cAMP (orange), and free *Sg* 2-MIBS (purple) are shown (left). Data points are shown as mean values. Error bars represent the standard deviation of three independent experiments. Values for $k_{cat}$, $K_m$, and $k_{cat}/K_m$ were derived by fitting the data to an allosteric sigmoidal curve in GraphPad Prism v 9.1.0, with the assumption that the calculated $K_{half}$ value is equivalent to $K_m$ (right).

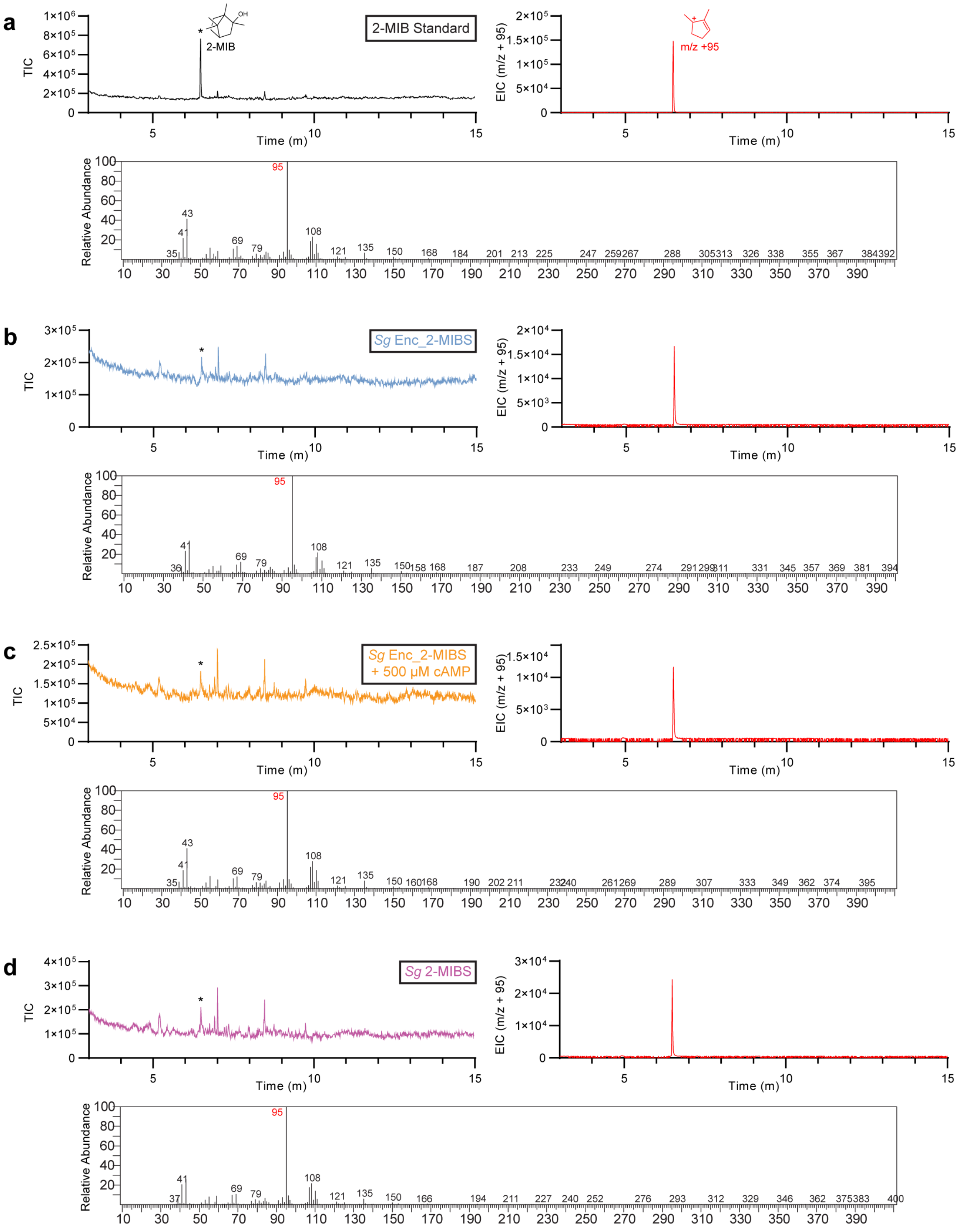


**Supplementary Fig. 13. Additional GC-MS data for *Sg* Enc_2-MIBS, *Sg* Enc, and 2-MIBS product profile assays. a**, GC-MS data for the 2-MIB standard. Total ion count (TIC) (top left); extracted ion chromatogram (EIC) (top right), and mass spectrum of the 2-MIB peak (bottom) are shown. The 2-MIB peak on the total ion current chromatogram is represented by the * symbol. **b**, GC-MS data for *Sg* Enc_2-MIBS. **c**, GC-MS data for *Sg* Enc_2-MIBS + 500 µM cAMP. **d**, GC-MS data for free *Sg* 2-MIBS.

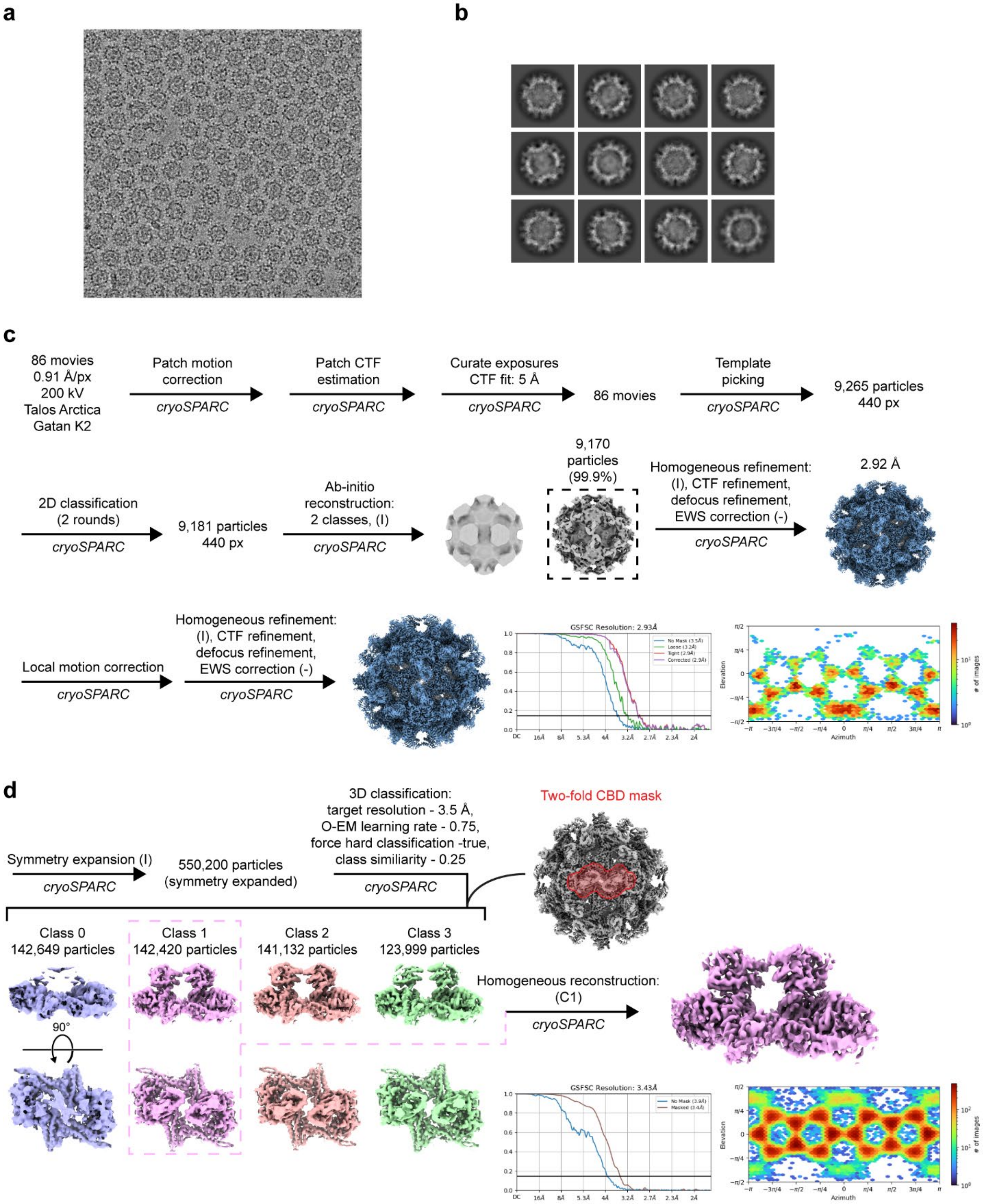


**Supplementary Fig. 14. Cryo-EM workflow for *Sg* Enc_2-MIBS + 20 mM cAMP. a**, Representative cryo-EM micrograph. **b**, Representative 2D class averages. **c**, Cryo-EM workflow. **d**, 3D classification workflow.

### Supplementary Table 1. Cryo-EM data collection, refinement and validation statistics.

| | *Sg* Enc_2-MIBS (*S. coelicolor* A3(2)-expressed) (EMDB-44554) (PDB ID: 9BHV) | *Sg* Enc (*E. coli*-expressed) (EMDB-44553) (PDB ID: 9BHU) | *Sg* Enc (*S. coelicolor* A3(2)-expressed) | *Sg*_Enc_2-MIBS + 20 mM cAMP (EMDB-44557) (PDB ID: 9BI0) |
|---|---|---|---|---|
| **Data collection and processing** | | | | |
| Magnification | 45,000x | 45,000x | 45,000x | 45,000x |
| Voltage (kV) | 200 | 200 | 200 | 200 |
| Electron exposure (e–/Å$^2$) | 42.94 | 43.22 | 37.95 | 39.98 |
| Defocus range (μm) | -0.8 to -1.8 | -1.0 to -1.8 | -1.0 to -1.8 | -0.8 to -1.8 |
| Pixel size (Å) | 0.91 | 0.91 | 0.91 | 0.91 |
| Symmetry imposed | I | I | I | I |
| Initial particle images (no.) | 66,966 | 159,394 | 7,003 | 9,265 |
| Final particle images (no.) | 51,173 | 135,027 | 4,091 | 9,170 |
| Map resolution (Å) | 3.02 | 2.71 | 3.47 | 2.93 |
| FSC threshold | 0.143 | 0.143 | 0.143 | 0.143 |
| **Refinement** | | | | |
| Initial model used (PDB code) | AlphaFold | AlphaFold | - | AlphaFold |
| Model resolution (Å) | 3.4 | 3.1 | - | 3.4 |
| FSC threshold | 0.5 | 0.5 | - | 0.5 |
| Map sharpening *B* factor (Å$^2$) | -115.8 | -127.9 | -81.0 | -92.0 |
| Model composition | | | | |
| Non-hydrogen | 3,099 | 3,099 | - | 3,099 |
| atoms | 395 | 395 | - | 395 |
| Protein residues | 0 | 0 | - | 0 |
| Ligands | | | | |
| *B* factors (Å$^2$) | | | | |
| Protein | 78.58 | 46.1 | - | 58.36 |
| Ligands | - | - | - | - |
| R.m.s. deviations | | | | |
| Bond lengths (Å) | 0.005 | 0.010 | - | 0.004 |
| Bond angles (°) | 1.016 | 1.062 | - | 0.985 |
| Validation | | | | |
| MolProbity score | 1.72 | 1.65 | - | 1.61 |
| Clashscore | 9.74 | 7.46 | - | 7.14 |
| Poor rotamers (%) | 0.61 | 0.92 | - | 0.92 |
| Ramachandran plot | | | | |
| Favored (%) | 96.66 | 96.40 | - | 96.66 |
| Allowed (%) | 3.34 | 3.6 | - | 3.34 |
| Disallowed (%) | 0 | 0 | - | 0 |

**Supplementary Table 2. Protein sequences of constructs used in this study.**

| Name | Protein Sequence |
|---|---|
| *Sg* Enc_2-MIBS | MTVDSTSEARLEVPRQSSLGTAAARNLASTTKSAPQMQEITSRWL<br>LRMLPWVETKGGAYRVNRRLTFTVGDGRVEFVQDGSTVRVIPQEL<br>GELALLRDFDDAEVLSAIADRCVQRDFRAGETLVERGTAADELHLIA<br>HGRIGQASAGSYGDEVTLDVLADGDRFGEHALLDEDARWSQTATA<br>ETSGTLLTLSRADFAAVVANSPALRSHLAAFTARSEQRQNHRGEAEI<br>AMSAGHVGEHELPGAFADYELKPREYELSVAQTILRVHTRVADLYN<br>GPMNQTEEQLRLTIEALRERQEHELINNREFGLLHNADFKQRIQTH<br>SGPPTPDDLDELLCRRRGTKFFLAHPRTIAAMGREFNARGLYPDHT<br>DLGGQQVPAWRGVPILPCGKIPITPERTSSILALRTGEEDQGVIGLR<br>QTGLPDEYEPGLSVRFMNIDEKAIISYLVSTYYSAAILVPDAVGVLEN<br>VQIANWPR* |
| | MPVPELPPPRSSLPEAVTRFGASVLGAVAARAHDSEATVGGPSGG<br>RPLPSPPAGLSFGPPSPAAPSADVPAPEAPGRGADLERLLCGPHG<br>LGTAGLRLTPGKERPVPATAREGRPIPGLYHHPVPEPDEARVEEVS<br>RRIKAWALDEVSLYPEEWEEQFDGFSVGRYMVGCHPDAPTVDHL<br>MLATRLMVAENAVDDCYCEDHGGSPVGLGERLLLAHTALDPLYTAR<br>EYQPGWAASLHADAPRRAYRSAMDYFVRAAGPSQADRLRHDMA<br>RLHLGYLAEAAWAQQDQVPEVWEYLAMRQFNNFRPCPTITDTVG<br>GYELPADLHAQAAMQKVIALASNATTIVNDLYSYTKELAAPGRHLNL<br>PVVIAEREGLSDQDAYLKSVEIHNELMHAFESEAAALAAACPVPSV<br>QRFLRGVAAWVDGNHHWHRSNTYRYSLPDFW* |
| *Sg* Enc | MTVDSTSEARLEVPRQSSLGTAAARNLASTTKSAPQMQEITSRWL<br>LRMLPWVETKGGAYRVNRRLTFTVGDGRVEFVQDGSTVRVIPQEL<br>GELALLRDFDDAEVLSAIADRCVQRDFRAGETLVERGTAADELHLIA<br>HGRIGQASAGSYGDEVTLDVLADGDRFGEHALLDEDARWSQTATA<br>ETSGTLLTLSRADFAAVVANSPALRSHLAAFTARSEQRQNHRGEAEI<br>AMSAGHVGEHELPGAFADYELKPREYELSVAQTILRVHTRVADLYN<br>GPMNQTEEQLRLTIEALRERQEHELINNREFGLLHNADFKQRIQTH<br>SGPPTPDDLDELLCRRRGTKFFLAHPRTIAAMGREFNARGLYPDHT<br>DLGGQQVPAWRGVPILPCGKIPITPERTSSILALRTGEEDQGVIGLR<br>QTGLPDEYEPGLSVRFMNIDEKAIISYLVSTYYSAAILVPDAVGVLEN<br>VQIANWPR* |
| *Sg* Enc_2-MIBS(ΔCLD) | MTVDSTSEARLEVPRQSSLGTAAARNLASTTKSAPQMQEITSRWL<br>LRMLPWVETKGGAYRVNRRLTFTVGDGRVEFVQDGSTVRVIPQEL<br>GELALLRDFDDAEVLSAIADRCVQRDFRAGETLVERGTAADELHLIA<br>HGRIGQASAGSYGDEVTLDVLADGDRFGEHALLDEDARWSQTATA<br>ETSGTLLTLSRADFAAVVANSPALRSHLAAFTARSEQRQNHRGEAE<br>IAMSAGHVGEHELPGAFADYELKPREYELSVAQTILRVHTRVADLY<br>NGPMNQTEEQLRLTIEALRERQEHELINNREFGLLHNADFKQRIQT<br>HSGPPTPDDLDELLCRRRGTKFFLAHPRTIAAMGREFNARGLYPD<br>HTDLGGQQVPAWRGVPILPCGKIPITPERTSSILALRTGEEDQGVIG<br>LRQTGLPDEYEPGLSVRFMNIDEKAIISYLVSTYYSAAILVPDAVGVL<br>ENVQIANWPR* |

MGRPIPGLYHHPVPEPDEARVEEVSRRIKAWALDEVSLYPEEWEE
QFDGFSVGRYMVGCHPDAPTVDHLMLATRLMVAENAVDDCYCED
HGGSPVGLGERLLLAHTALDPLYTAREYQPGWAASLHADAPRRAY
RSAMDYFVRAAGPSQADRLRHDMARLHLGYLAEAAWAQQDQVP
EVWEYLAMRQFNNFRPCPTITDTVGGYELPADLHAQAAMQKVIAL
ASNATTIVNDLYSYTKELAAPGRHLNLPVVIAEREGLSDQDAYLKSV
EIHNELMHAFESEAAALAAACPVPSVQRFLRGVAAWVDGNHHWH
RSNTYRYSLPDFW*

| | |
|---|---|
| *Sg* Enc_CLD-mNeonGreen | MTVDSTSEARLEVPRQSSLGTAAARNLASTTKSAPQMQEITSRWL LRMLPWVETKGGAYRVNRRLTFTVGDGRVEFVQDGSTVRVIPQEL GELALLRDFDDAEVLSAIADRCVQRDFRAGETLVERGTAADELHLIA HGRIGQASAGSYGDEVTLDVLADGDRFGEHALLDEDARWSQTATA ETSGTLLTLSRADFAAVVANSPALRSHLAAFTARSEQRQNHRGEAE IAMSAGHVGEHELPGAFADYELKPREYELSVAQTILRVHTRVADLY NGPMNQTEEQLRLTIEALRERQEHELINNREFGLLHNADFKQRIQT HSGPPTPDDLDELLCRRRGTKFFLAHPRTIAAMGREFNARGLYPD HTDLGGQQVPAWRGVPILPCGKIPITPERTSSILALRTGEEDQGVIG LRQTGLPDEYEPGLSVRFMNIDEKAIISYLVSTYYSAAILVPDAVGVL ENVQIANWPR* |
| | MPVPELPPPRSSLPEAVTRFGASVLGAVAARAHDSEATVGGPSGG RPLPSPPAGLSFGPPSPAAPSADVPAPEAPGRGADLERLLCGPHG LGTAGLRLTPGKERPVPATAREGGSVSKGEEDNMASLPATHELHIF GSINGVDFDMVGQGTGNPNDGYEELNLKSTKGDLQFSPWILVPHI GYGFHQYLPYPDGMSPFQAAMVDGSGYQVHRTMQFEDGASLTV NYRYTYEGSHIKGEAQVKGTGFPADGPVMTNSLTAADWCRSKKTY PNDKTIISTFKWSYTTGNGKRYRSTARTTYTFAKPMAANYLKNQPM YVFRKTELKHSKTELNFKEWQKAFTDVMGMDELYK* |
| *Sg* Enc_CLD-mNeonGreen (ΔCLD_B) | MTVDSTSEARLEVPRQSSLGTAAARNLASTTKSAPQMQEITSRWL LRMLPWVETKGGAYRVNRRLTFTVGDGRVEFVQDGSTVRVIPQEL GELALLRDFDDAEVLSAIADRCVQRDFRAGETLVERGTAADELHLIA HGRIGQASAGSYGDEVTLDVLADGDRFGEHALLDEDARWSQTATA ETSGTLLTLSRADFAAVVANSPALRSHLAAFTARSEQRQNHRGEAE IAMSAGHVGEHELPGAFADYELKPREYELSVAQTILRVHTRVADLY NGPMNQTEEQLRLTIEALRERQEHELINNREFGLLHNADFKQRIQT HSGPPTPDDLDELLCRRRGTKFFLAHPRTIAAMGREFNARGLYPD HTDLGGQQVPAWRGVPILPCGKIPITPERTSSILALRTGEEDQGVIG LRQTGLPDEYEPGLSVRFMNIDEKAIISYLVSTYYSAAILVPDAVGVL ENVQIANWPR* |
| | MPVPELPPPRSSGGPSGGRPLPSPPAGLSFGPPSPAAPSADVPAP EAPGRGADLERLLCGPHGLGTAGLRLTPGKERPVPATAREGGSVS KGEEDNMASLPATHELHIFGSINGVDFDMVGQGTGNPNDGYEELN LKSTKGDLQFSPWILVPHIGYGFHQYLPYPDGMSPFQAAMVDGSG YQVHRTMQFEDGASLTVNYRYTYEGSHIKGEAQVKGTGFPADGPV MTNSLTAADWCRSKKTYPNDKTIISTFKWSYTTGNGKRYRSTARTT |

| | |
|---|---|
| | YTFAKPMAANYLKNQPMYVFRKTELKHSKTELNFKEWQKAFTDVM<br>GMDELYK* |
| *Sg* Enc_CLD-mNeonGreen (ΔCLD_D) | MTVDSTSEARLEVPRQSSLGTAAARNLASTTKSAPQMQEITSRWL<br>LRMLPWVETKGGAYRVNRRLTFTVGDGRVEFVQDGSTVRVIPQEL<br>GELALLRDFDDAEVLSAIADRCVQRDFRAGETLVERGTAADELHLIA<br>HGRIGQASAGSYGDEVTLDVLADGDRFGEHALLDEDARWSQTATA<br>ETSGTLLTLSRADFAAVVANSPALRSHLAAFTARSEQRQNHRGEAE<br>IAMSAGHVGEHELPGAFADYELKPREYELSVAQTILRVHTRVADLY<br>NGPMNQTEEQLRLTIEALRERQEHELINNREFGLLHNADFKQRIQT<br>HSGPPTPDDLDELLCRRRGTKFFLAHPRTIAAMGREFNARGLYPD<br>HTDLGGQQVPAWRGVPILPCGKIPITPERTSSILALRTGEEDQGVIG<br>LRQTGLPDEYEPGLSVRFMNIDEKAIISYLVSTYYSAAILVPDAVGVL<br>ENVQIANWPR* |
| | MPVPELPPPRSSLPEAVTRFGASVLGAVAARAHDSEATVGGPSGG<br>RPLPSPPAGLSFGPPSPAAPSADVPAPEAPGRGADPGKERPVPAT<br>AREGGSVSKGEEDNMASLPATHELHIFGSINGVDFDMVGQGTGNP<br>NDGYEELNLKSTKGDLQFSPWILVPHIGYGFHQYLPYPDGMSPFQ<br>AAMVDGSGYQVHRTMQFEDGASLTVNYRYTYEGSHIKGEAQVKG<br>TGFPADGPVMTNSLTAADWCRSKKTYPNDKTIISTFKWSYTTGNG<br>KRYRSTARTTYTFAKPMAANYLKNQPMYVFRKTELKHSKTELNFKE<br>WQKAFTDVMGMDELYK* |
| *Sg* Enc_CLD-mNeonGreen (ΔCLD_B,D) | MTVDSTSEARLEVPRQSSLGTAAARNLASTTKSAPQMQEITSRWL<br>LRMLPWVETKGGAYRVNRRLTFTVGDGRVEFVQDGSTVRVIPQEL<br>GELALLRDFDDAEVLSAIADRCVQRDFRAGETLVERGTAADELHLIA<br>HGRIGQASAGSYGDEVTLDVLADGDRFGEHALLDEDARWSQTATA<br>ETSGTLLTLSRADFAAVVANSPALRSHLAAFTARSEQRQNHRGEAE<br>IAMSAGHVGEHELPGAFADYELKPREYELSVAQTILRVHTRVADLY<br>NGPMNQTEEQLRLTIEALRERQEHELINNREFGLLHNADFKQRIQT<br>HSGPPTPDDLDELLCRRRGTKFFLAHPRTIAAMGREFNARGLYPD<br>HTDLGGQQVPAWRGVPILPCGKIPITPERTSSILALRTGEEDQGVIG<br>LRQTGLPDEYEPGLSVRFMNIDEKAIISYLVSTYYSAAILVPDAVGVL<br>ENVQIANWPR* |
| | MPVPELPPPRSSGGPSGGRPLPSPPAGLSFGPPSPAAPSADVPAP<br>EAPGRGADPGKERPVPATAREGGSVSKGEEDNMASLPATHELHIF<br>GSINGVDFDMVGQGTGNPNDGYEELNLKSTKGDLQFSPWILVPHI<br>GYGFHQYLPYPDGMSPFQAAMVDGSGYQVHRTMQFEDGASLTV<br>NYRYTYEGSHIKGEAQVKGTGFPADGPVMTNSLTAADWCRSKKTY<br>PNDKTIISTFKWSYTTGNGKRYRSTARTTYTFAKPMAANYLKNQPM<br>YVFRKTELKHSKTELNFKEWQKAFTDVMGMDELYK* |
| *Sg* Enc_CLD-mNeonGreen (ΔCLD_B,C,D,E) | MTVDSTSEARLEVPRQSSLGTAAARNLASTTKSAPQMQEITSRWL<br>LRMLPWVETKGGAYRVNRRLTFTVGDGRVEFVQDGSTVRVIPQEL<br>GELALLRDFDDAEVLSAIADRCVQRDFRAGETLVERGTAADELHLIA<br>HGRIGQASAGSYGDEVTLDVLADGDRFGEHALLDEDARWSQTATA<br>ETSGTLLTLSRADFAAVVANSPALRSHLAAFTARSEQRQNHRGEAE<br>IAMSAGHVGEHELPGAFADYELKPREYELSVAQTILRVHTRVADLY |

| | |
|---|---|
| | NGPMNQTEEQLRLTIEALRERQEHELINNREFGLLHNADFKQRIQT<br>HSGPPTPDDLDELLCRRRGTKFFLAHPRTIAAMGREFNARGLYPD<br>HTDLGGQQVPAWRGVPILPCGKIPITPERTSSILALRTGEEDQGVIG<br>LRQTGLPDEYEPGLSVRFMNIDEKAIISYLVSTYYSAAILVPDAVGVL<br>ENVQIANWPR* |
| | MPVPELPPPRSSGGSVSKGEEDNMASLPATHELHIFGSINGVDFD<br>MVGQGTGNPNDGYEELNLKSTKGDLQFSPWILVPHIGYGFHQYLP<br>YPDGMSPFQAAMVDGSGYQVHRTMQFEDGASLTVNYRYTYEGS<br>HIKGEAQVKGTGFPADGPVMTNSLTAADWCRSKKTYPNDKTIISTF<br>KWSYTTGNGKRYRSTARTTYTFAKPMAANYLKNQPMYVFRKTELK<br>HSKTELNFKEWQKAFTDVMGMDELYK* |
| *Sg* Enc_CLD-<br>mNeonGreen<br>(ΔCLD_A,B,D,E) | MTVDSTSEARLEVPRQSSLGTAAARNLASTTKSAPQMQEITSRWL<br>LRMLPWVETKGGAYRVNRRLTFTVGDGRVEFVQDGSTVRVIPQEL<br>GELALLRDFDDAEVLSAIADRCVQRDFRAGETLVERGTAADELHLIA<br>HGRIGQASAGSYGDEVTLDVLADGDRFGEHALLDEDARWSQTATA<br>ETSGTLLTLSRADFAAVVANSPALRSHLAAFTARSEQRQNHRGEAE<br>IAMSAGHVGEHELPGAFADYELKPREYELSVAQTILRVHTRVADLY<br>NGPMNQTEEQLRLTIEALRERQEHELINNREFGLLHNADFKQRIQT<br>HSGPPTPDDLDELLCRRRGTKFFLAHPRTIAAMGREFNARGLYPD<br>HTDLGGQQVPAWRGVPILPCGKIPITPERTSSILALRTGEEDQGVIG<br>LRQTGLPDEYEPGLSVRFMNIDEKAIISYLVSTYYSAAILVPDAVGVL<br>ENVQIANWPR* |
| | MGGPSGGRPLPSPPAGLSFGPPSPAAPSADVPAPEAPGRGADGG<br>SVSKGEEDNMASLPATHELHIFGSINGVDFDMVGQGTGNPNDGYE<br>ELNLKSTKGDLQFSPWILVPHIGYGFHQYLPYPDGMSPFQAAMVD<br>GSGYQVHRTMQFEDGASLTVNYRYTYEGSHIKGEAQVKGTGFPA<br>DGPVMTNSLTAADWCRSKKTYPNDKTIISTFKWSYTTGNGKRYRS<br>TARTTYTFAKPMAANYLKNQPMYVFRKTELKHSKTELNFKEWQKA<br>FTDVMGMDELYK* |
| *Sg* Enc_CLD-<br>mNeonGreen<br>(ΔCLD_A,B,C) | MTVDSTSEARLEVPRQSSLGTAAARNLASTTKSAPQMQEITSRWL<br>LRMLPWVETKGGAYRVNRRLTFTVGDGRVEFVQDGSTVRVIPQEL<br>GELALLRDFDDAEVLSAIADRCVQRDFRAGETLVERGTAADELHLIA<br>HGRIGQASAGSYGDEVTLDVLADGDRFGEHALLDEDARWSQTATA<br>ETSGTLLTLSRADFAAVVANSPALRSHLAAFTARSEQRQNHRGEAE<br>IAMSAGHVGEHELPGAFADYELKPREYELSVAQTILRVHTRVADLY<br>NGPMNQTEEQLRLTIEALRERQEHELINNREFGLLHNADFKQRIQT<br>HSGPPTPDDLDELLCRRRGTKFFLAHPRTIAAMGREFNARGLYPD<br>HTDLGGQQVPAWRGVPILPCGKIPITPERTSSILALRTGEEDQGVIG<br>LRQTGLPDEYEPGLSVRFMNIDEKAIISYLVSTYYSAAILVPDAVGVL<br>ENVQIANWPR* |
| | MLERLLCGPHGLGTAGLRLTPGKERPVPATAREGGSVSKGEEDNM<br>ASLPATHELHIFGSINGVDFDMVGQGTGNPNDGYEELNLKSTKGDL<br>QFSPWILVPHIGYGFHQYLPYPDGMSPFQAAMVDGSGYQVHRTM<br>QFEDGASLTVNYRYTYEGSHIKGEAQVKGTGFPADGPVMTNSLTA<br>ADWCRSKKTYPNDKTIISTFKWSYTTGNGKRYRSTARTTYTFAKPM |

| | |
|---|---|
| | AANYLKNQPMYVFRKTELKHSKTELNFKEWQKAFTDVMGMDELY<br>K* |
| *Sg* Enc_CLD-mNeonGreen (ΔCLD_A,B,C,D) | MTVDSTSEARLEVPRQSSLGTAAARNLASTTKSAPQMQEITSRWL<br>LRMLPWVETKGGAYRVNRRLTFTVGDGRVEFVQDGSTVRVIPQEL<br>GELALLRDFDDAEVLSAIADRCVQRDFRAGETLVERGTAADELHLIA<br>HGRIGQASAGSYGDEVTLDVLADGDRFGEHALLDEDARWSQTATA<br>ETSGTLLTLSRADFAAVVANSPALRSHLAAFTARSEQRQNHRGEAE<br>IAMSAGHVGEHELPGAFADYELKPREYELSVAQTILRVHTRVADLY<br>NGPMNQTEEQLRLTIEALRERQEHELINNREFGLLHNADFKQRIQT<br>HSGPPTPDDLDELLCRRRGTKFFLAHPRTIAAMGREFNARGLYPD<br>HTDLGGQQVPAWRGVPILPCGKIPITPERTSSILALRTGEEDQGVIG<br>LRQTGLPDEYEPGLSVRFMNIDEKAIISYLVSTYYSAAILVPDAVGVL<br>ENVQIANWPR* |
| | MPGKERPVPATAREGGSVSKGEEDNMASLPATHELHIFGSINGVDF<br>DMVGQGTGNPNDGYEELNLKSTKGDLQFSPWILVPHIGYGFHQYL<br>PYPDGMSPFQAAMVDGSGYQVHRTMQFEDGASLTVNYRYTYEG<br>SHIKGEAQVKGTGFPADGPVMTNSLTAADWCRSKKTYPNDKTIIST<br>FKWSYTTGNGKRYRSTARTTYTFAKPMAANYLKNQPMYVFRKTEL<br>KHSKTELNFKEWQKAFTDVMGMDELYK* |
| *Sg* Enc_CLD-mNeonGreen (ΔCLD) | MTVDSTSEARLEVPRQSSLGTAAARNLASTTKSAPQMQEITSRWL<br>LRMLPWVETKGGAYRVNRRLTFTVGDGRVEFVQDGSTVRVIPQEL<br>GELALLRDFDDAEVLSAIADRCVQRDFRAGETLVERGTAADELHLIA<br>HGRIGQASAGSYGDEVTLDVLADGDRFGEHALLDEDARWSQTATA<br>ETSGTLLTLSRADFAAVVANSPALRSHLAAFTARSEQRQNHRGEAE<br>IAMSAGHVGEHELPGAFADYELKPREYELSVAQTILRVHTRVADLY<br>NGPMNQTEEQLRLTIEALRERQEHELINNREFGLLHNADFKQRIQT<br>HSGPPTPDDLDELLCRRRGTKFFLAHPRTIAAMGREFNARGLYPD<br>HTDLGGQQVPAWRGVPILPCGKIPITPERTSSILALRTGEEDQGVIG<br>LRQTGLPDEYEPGLSVRFMNIDEKAIISYLVSTYYSAAILVPDAVGVL<br>ENVQIANWPR* |
| | MVSKGEEDNMASLPATHELHIFGSINGVDFDMVGQGTGNPNDGYE<br>ELNLKSTKGDLQFSPWILVPHIGYGFHQYLPYPDGMSPFQAAMVD<br>GSGYQVHRTMQFEDGASLTVNYRYTYEGSHIKGEAQVKGTGFPA<br>DGPVMTNSLTAADWCRSKKTYPNDKTIISTFKWSYTTGNGKRYRS<br>TARTTYTFAKPMAANYLKNQPMYVFRKTELKHSKTELNFKEWQKA<br>FTDVMGMDELYK* |
| *Sg* 2-MIBS | MPVPELPPPRSSLPEAVTRFGASVLGAVAARAHDSEATVGGPSGG<br>RPLPSPPAGLSFGPPSPAAPSADVPAPEAPGRGADLERLLCGPHG<br>LGTAGLRLTPGKERPVPATAREGRPIPGLYHHPVPEPDEARVEEVS<br>RRIKAWALDEVSLYPEEWEEQFDGFSVGRYMVGCHPDAPTVDHL<br>MLATRLMVAENAVDDCYCEDHGGSPVGLGERLLLAHTALDPLYTA<br>REYQPGWAASLHADAPRRAYRSAMDYFVRAAGPSQADRLRHDM<br>ARLHLGYLAEAAWAQQDQVPEVWEYLAMRQFNNFRPCPTITDTV<br>GGYELPADLHAQAAMQKVIALASNATTIVNDLYSYTKELAAPGRHL<br>NLPVVIAEREGLSDQDAYLKSVEIHNELMHAFESEAAALAAACPVP |

| | |
|---|---|
| | SVQRFLRGVAAWVDGNHHWHRSNTYRYSLPDFWS*ENLYFQG*GS<br>GHHHHHHHH* |
| *Sg* MT | MHHHHHHHHGGGSGGGS*ENLYFQG*MTNAELTTAPTLRIPGPATPY<br>QGDIARYWDGEARPVNLRLGDVDGLYHHHYGIGEVDTASLGNPED<br>SESEKKLITELHRLESAQADFLLGHLGDIGRDDTLVDAGCGRGGSM<br>VMAHQRFGCSVEGVTLSAKQADFANGRAAELGIGDHVRARVCNM<br>LSTPFATGSAAASWNNESSMYVDLDDLFAEHSRVLKVGGRYVTIT<br>GCWNPRYGQPSKWVSQINAHFECNIHSRREYLRAMADNRLVPQA<br>VIDLTPDTLPYWELRATSSLVTGIEDAFINSYRDGSFQYLLIAADRV* |
| *Ec* CAP | MHHHHHH*ENLYFQG*GGSGMVLGKPQTDPTLEWFLSHCHIHKYPS<br>KSTLIHQGEKAETLYYIVKGSVAVLIKDEEGKEMILSYLNQGDFIGEL<br>GLFEEGQERSAWVRAKTACEVAEISYKKFRQLIQVNPDILMRLSAQ<br>MARRLQVTSEKVGNLAFLDVTGRIAQTLLNLAKQPDAMTHPDGMQ<br>IKITRQEIGQIVGCSRETVGRILKMLEDQNLISAHGKTIVVYGTR* |

[1]Stops are represented by asterisks

[2]Underlined amino acids represent expression tags

[3]Italicized amino acids represent TEV protease cut sites

### Supplementary References


1 Benisch, R., Andreas, M. P. & Giessen, T. W. A widespread bacterial protein compartment sequesters and stores elemental sulfur. *Science Advances* **10**, eadk9345, doi:doi:10.1126/sciadv.adk9345 (2024).

2 Nichols, R. J. *et al.* Discovery and characterization of a novel family of prokaryotic nanocompartments involved in sulfur metabolism. *eLife* **10**, doi:10.7554/elife.59288 (2021).

3 Jumper, J. *et al.* Highly accurate protein structure prediction with AlphaFold. *Nature* **596**, 583-589, doi:10.1038/s41586-021-03819-2 (2021).

4 Ashkenazy, H. *et al.* ConSurf 2016: an improved methodology to estimate and visualize evolutionary conservation in macromolecules. *Nucleic Acids Res* **44**, W344-350, doi:10.1093/nar/gkw408 (2016).
